# Optimizing Intermediate Representations: A Framework for Low-Cost, High-Accuracy Behavior Quantification

**DOI:** 10.64898/2026.03.29.715170

**Authors:** Jessica D. Choi, Brian Geuther, Vivek Kumar

## Abstract

Quantitative measurement of animal behavior is a cornerstone of neuroscience, genetics, and ethology. While modern computer vision has democratized automated analysis, the field has coalesced around pose estimation as the standard intermediate representation. This reliance imposes a significant bottleneck: researchers must often train custom pose models using large, labor-intensive datasets. Furthermore, the assumption that denser anatomical tracking yields better classification remains largely unverified. Here, we benchmark intermediate representations for supervised mouse behavior classification to determine the optimal trade-off between annotation cost and model performance. We systematically evaluate the sensitivity of classification to keypoint density, the impact of temporal feature engineering, and the viability of segmentation-derived shape descriptors as a low-cost alternative. We find that classifier performance is remarkably robust to keypoint variation; increasing keypoint density yields negligible gains, particularly when behavior training sets are sufficiently large. In contrast, augmenting models with temporal features (specifically FFT-based signal processing) consistently drives performance improvements. Crucially, we demonstrate that whole-body segmentation achieves performance parity with explicit pose estimation across most behaviors. These findings challenge the "more is better" intuition in pose tracking and suggest a paradigm shift: efficient pipelines should prioritize behavioral dataset volume and temporal dynamics over complex anatomical keypoints.

## 1 Introduction

Quantitative analysis of animal behavior is a cornerstone of modern neurogenetics. However, the study of lab animal behavior has traditionally relied on manual scoring by behavioral experts with little automation. New analysis tools are often the driver of discoveries in behavioral biology by providing scalable, reproducible, and objective analyses. For example, the open field assay, when developed in 1934 relied entirely on manual scoring [1, 2]. One of its first automations was the integration of infrared beams to infer rodent position, followed by computer vision methods using background subtraction to track animals as a "blob". While effective, these early methods were limited to simple environments.

With the proliferation of deep learning technology, biologists now have access to a suite of tools designed to automatically extract meaningful, detailed measurements from these environments [3, 4, 5]. A major advance has been the application of neural networks for pose estimation, allowing the tracking of specific animal body parts (keypoints) with high accuracy. Tools such as DeepLabCut, SLEAP, DeepPoseKit, and APT have become standard for extracting kinematic data [6, 7, 8, 9]. This rich, high-dimensional pose data serves as an intermediate representation for downstream behavior classifiers, including supervised toolkits like MARS, SIMBA, A-SOiD, and JABS, as well as unsupervised methods like MotionMapper, Keypoint-MoSeq, B-SOiD, and VAME, and foundation model approaches like VideoPrism, V-JEPA 2, and FERAL [10, 11, 12, 5, 13, 14, 15, 16, 17, 18, 19].

This use of pose as an intermediate representation directly parallels human skeleton-based action recognition, a mature field in computer vision [20] predicated on the same principle: that skeletal joint kinematics provide a powerful, background-invariant representation for classifying behavior. The history of pose estimation predates its adoption in neuroscience by decades. Early markerless tracking relied on hand-crafted feature detectors with limited applicability to the varied conditions of laboratory recordings. The field was transformed when convolutional neural networks, having demonstrated their power on large annotated image datasets [21], were extended to pose estimation [22]. From there, the progression to neuroscience was direct: many of the pose estimation tools now standard in animal behavioral research are adaptations of human pose estimation architectures [23]. The field inherited powerful methods from computer vision, but with them came a critical challenge—these methods were developed and validated at a scale of annotation that animal behavioral research has not yet matched. For reference, the COCO keypoint benchmark alone contains over 250,000 annotated person instances across 200,000 images [24], and the MPII Human Pose dataset provides over 40,000 [25]. The largest comparable resource for animals, AP-10K, comprises just 10,015 images spanning 54 species [26], while most individual laboratory projects rely on only a few hundred labeled frames.

Therefore, despite the popularity and utility of pose estimation, its adoption for animals, particularly rodents, is fundamentally limited by the cost of generating annotated datasets. This creates a critical bottleneck, as researchers must invest significant annotation effort in two separate stages: (1) labeling keypoints to train the pose estimation model, and (2) labeling behavior bouts to train the downstream classifier. These costs are not symmetric. Keypoint annotation is slow, ranging from 0.90 to 1.77 seconds per keypoint [4]. In contrast, behavior annotation costs only 0.16 to 0.25 seconds per frame [4]. Furthermore, it is often unclear which keypoints are most informative, leading labs to define keypoint sets based on intuition, often assuming more is better [27]. This leads to a central, un-benchmarked question: what is the optimal trade-off between keypoint annotation and behavior annotation costs?

The effort required to build pose estimation models that generalize reliably adds to this cost. Even under a best case scenario, training an initial pose model requires at least 200 labeled frames [28], but this estimate only holds up when new videos very closely replicate conditions from training. This is an overfitting approach i.e., where the model is trained and inferred on the same specific setup. When recordings change (slightly different camera angle, lighting setup, environmental enrichment, animal size, or coat color), performance goes down and additional annotation is required to re-attain reasonable performance. This means that pose is not a one-time investment. Rather, it becomes more of an iterative annotation cycle, where researchers extract new frames from every new experiment, re-annotate keypoints, and retrain the model. This initially assumed low upfront annotation cost is not fixed once, instead for many groups it becomes a recurring price for every new experiment. Beyond this, data that has been previously inferred needs to be re-inferred with the new model. For example, if a researcher runs pose inference of 20 videos using one model, and then needs to add one more video to retrain, the user should then re-infer on all the videos to maintain model consistency across results.

To circumvent this cycle, the field has attempted to build models that can generalize across new data from the outset. For example, in our open field setup, we found that it required nearly 9000 annotated frames to generalize across genetic diversity, size, sex, and coat color [29]. This is an order of magnitude above standard recommendations. This shows a discrepancy between the definition of generalization that is typically used when evaluating pose models and broader generalization that is actually required in practice. The standard benchmark of 200 annotated frames can produce high accuracy on held-out frames extracted from the same recording session and conditions, which is measuring within-distribution performance rather than generalization across videos. Meanwhile, the human pose estimation field has state-of-the-art models trained on hundreds of thousands of annotations spanning diverse clothing, lighting, camera angles, and environments [24]. The human problem is admittedly harder, with much more variation in context compared to a controlled lab recording, but the principle is the same. For true generalization across multiple conditions, annotation costs scale accordingly.

Thus far, there has been no truly generalizable pose estimator that reliably works across novel laboratory setups without any extra annotation. In practice, many groups that are using novel arenas still require at least some targeted re-annotation before they can reach reliable predictions. This boundary of what is a "novel context" is not cleanly defined, but the fact that active refinement loops are recommended as part of the standard workflow reflects how far the field is from a truly annotation-free pose estimation.

Although keypoint-based representation has gained traction, it is not the only approach. Early classifiers like JAABA [30] operated effectively on features derived from ellipse-fitting and segmentation. This concept remains powerful; in previous work, modern segmentation-based features were used to accurately classify sleep states (REM, NREM, Wake) [31]. Such tasks are now even more feasible at scale due to foundational models like Segment Anything (SAM), [32], SAM2 [33], and SAM3 [34]. Alternatively, neural networks can classify behavior directly from raw pixels, as seen in DeepEthogram [35] and other custom CNNs. These end-to-end approaches mirror a trend in mainstream computer vision, exemplified by state-of-the-art video foundation models like VideoMAE [36], V-JEPA 2 [18], and FERAL [19]. However, these state-of-the-art models require massive training datasets to generalize. VideoMAE was pretrained on Kinetics-400, a dataset of approximately 240,000 video clips [36]; VideoPrism was trained on 36 million curated video-text pairs plus 582 million additional video clips [17]; and V-JEPA 2 was pretrained on over 1 million hours of internet video [18]. This volume of data is unattainable for most individual biology labs. For many scientific applications, intermediate representations remain the more practical path. Yet the question of which intermediate representation is best, and at what cost, has not been answered.

Despite the significant cost of keypoint annotation, two core assumptions underlying current pipelines remain untested. The first is that more keypoints yield better behavior classifiers. In practice, many groups default to large keypoint sets under the assumption that denser body coverage produces richer features, yet this relationship has never been systematically evaluated. The second, more fundamental assumption is that keypoints are necessary at all. As pose estimation tools became available, the field converged on keypoint-based classification without direct evidence of its superiority over alternatives. To our knowledge, no controlled comparison of keypoint-based and segmentation-based features for supervised behavior classification has been conducted. Both assumptions have shaped how the field allocates its most expensive resource—annotation effort—without empirical support.

In this paper we evaluate how keypoint selection affects performance across behaviors to understand the risk of information loss associated with sub-optimal keypoint selection. We provide evidence for the cost trade-offs associated with selecting intermediate representations, dataset annotation cost, and achieving desired classifier performance. We address three fundamental questions: (1) How much does the specific choice and number of keypoints matter for downstream classification? (2) How much do temporal features impact performance, and which features are best? (3) Are keypoints, with their high annotation cost, necessary at all, or can modern segmentation-based features achieve comparable performance? Our findings suggest that annotation effort is often better spent labeling more behaviors rather than more keypoints, and that for many applications, simpler, low-cost segmentation features are a competitive alternative.

## 2 Results

This work is motivated by the asymmetry present in annotation costs between the two stages of building a behavior classifier. Keypoint annotation costs 0.90 to 1.77 seconds per keypoint [4]. Previous work from our group required 8910 frames of 12 keypoints in our JABS framework [5] to develop a generalizable model [29]. At the low estimate of 0.9 seconds per keypoint, this dataset would have taken 26.7 hours to annotate, and at the high end of 1.77 seconds, it would have taken 52.6 hours. There is a motivation to reduce the number of keypoints annotated to save significantly on time-cost in annotating keypoints. In the next stage of classifier building, behavior annotation costs 0.16-0.25 seconds per frame [4]. To annotate 100,000 behavior frames (which is an order of magnitude more than most released behavior annotation sets, Table S6), it would only cost 4.44-6.94 hours to annotate. We do recommend a targeted strategy of annotation, focusing on borderline predictions [5], which may also produce higher-quality classifiers in a annotation-efficient manner. With these costs in mind, it is important to determine where the time-cost and efforts are most valuable for the best performance.

### 2.1 Behavior Classification Performance is Robust to Variation in Literature-based Keypoint Sets

A primary bottleneck in modern behavioral quantification is the manual labor required to annotate datasets. Researchers must often decide how many body parts (keypoints) to track, frequently operating under the intuitive assumption that maximizing anatomical detail is a prerequisite for high-accuracy behavior classification. However, this relationship between pose complexity and classifier performance has not been systematically tested. We sought to determine if dense keypoint annotation is strictly necessary, or if streamlined representations could achieve comparable accuracy, thereby reducing the burden on experimentalists. If keypoint selection and quantity meaningfully impact classifier performance, we would expect to see performance differences between sets with more keypoints consistently improving performance.

To systematically evaluate these questions, we compared downstream classifier performance across four standard keypoint sets established in the literature (JABS [5], MARS [10], MoSeq [37], and Mouse Resource [14]). These sets vary two-fold in keypoint number (5 to 12), and four-fold in feature count (462 to 1947). These sets are described in Table 1, which shows the specific keypoints included in each configuration as well as the resulting number of features we computed from each set (red points indicate the keypoints present, whereas the gray points indicate excluded keypoints).

**Table 1:**
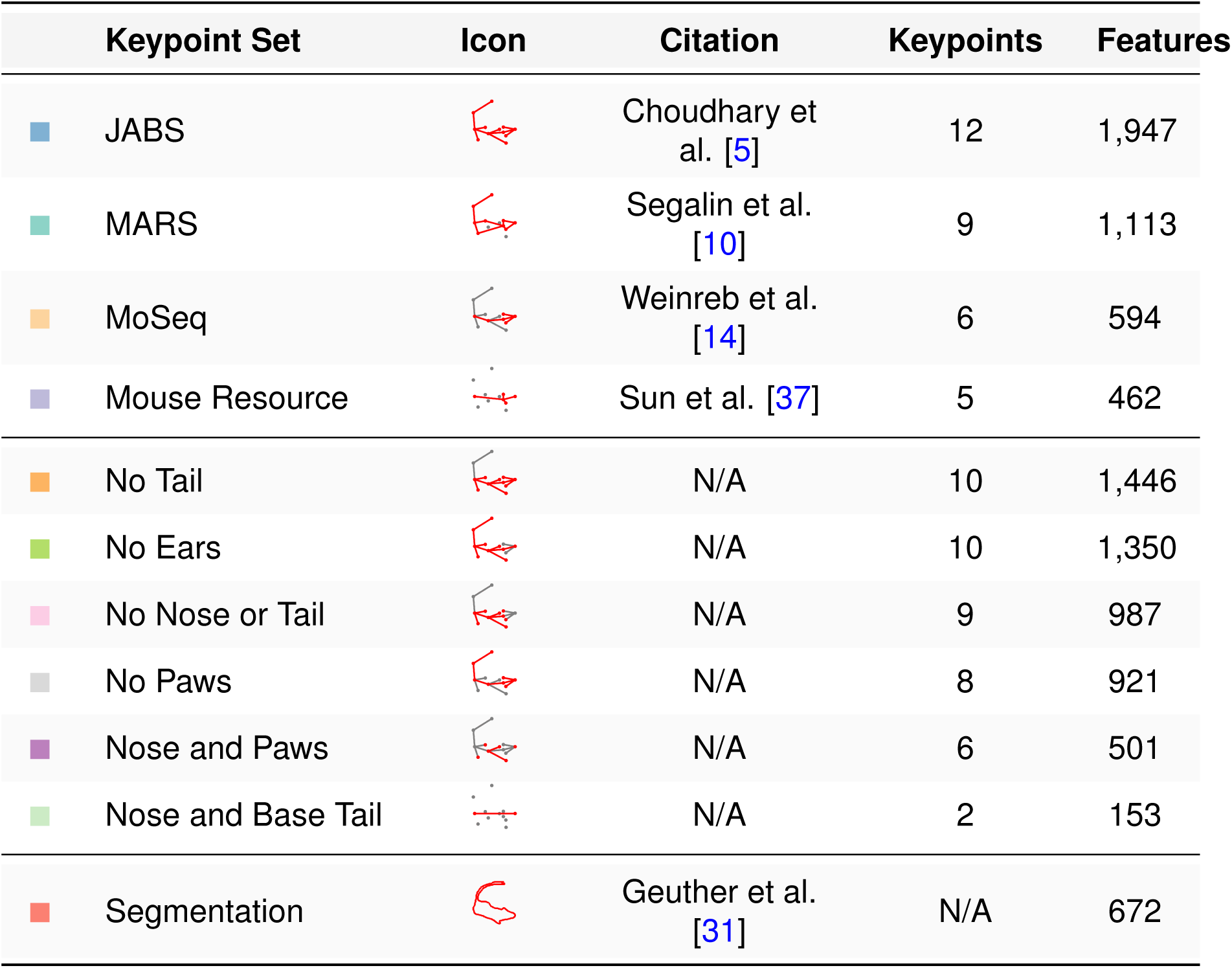
Keypoint sets evaluated for behavior classifier performance. (Top rows) Literature-based keypoint sets from published behavior analysis tools. (Middle rows) Ablations tested to determine impact of removing specific body parts. (Bottom row) Segmentation-based features tested that provide an alternative pose-free representation. The icons show the tracked keypoints for each set (red indicates presence, gray indicates absence). Each set of keypoints has features calculated which are not scaled linearly to keypoint number, as most features are computed from keypoint combinations.

For all comparisons in the section, we computed features from keypoints using a combined temporal feature set (JABS and FFT window features, described in detail in Section 2.3). This feature set combines spatial information such as distances and angles (base features) across time windows to capture behavioral dynamics. We use this feature computation method across all keypoint sets to isolate the effect of keypoint selection from specific feature engineering choices. Our goal is to determine whether the choice of which body parts to track, or the number of keypoints affects performance.

Rigorous benchmarking of feature design ideally requires large, expertly labeled datasets. However, obtaining these labeled datasets remains a significant challenge in the animal behavior field. Labeling requires expert annotators, which is associated with high time-costs and requires an understanding of the nuance of each behavior. Many datasets contain only intermediate representations or behavior annotations, but finding a combination of both is challenging. To work within these constraints, we assembled a dataset spanning diverse behaviors by combining our own single-mouse data with other published datasets. Here we use rearing, scratching, turning, and grooming from our previously published work (Table S6, [38, 5]) alongside rearing behaviors from Sturman et al. [4]. We selected these behaviors as they represent varied types of behavior and movement, and importantly had high-quality ground truth labels available. We included two independent rearing datasets (JABS and Sturman) to evaluate keypoint set robustness across different annotation sources. The amount of available labeled data varies across our behaviors (Table S6), which further reflects the practical constraints of behavior annotation. While larger or more numerous test sets would be ideal, these datasets represent the best available ground truth for rigorous evaluation of keypoint set design.

We quantified performance using F1, which balances precision and recall [39]. For each behavior and keypoint set, we measured performance through cross-validation across videos (12-20 folds depending on behavior). For grooming evaluation we instead used a fixed heldout set from [38], therefore error bars are not applicable to this dataset. We ran statistical comparisons between the keypoint sets using the Friedman test [40, 41], a non-parametric repeated measures test that accounts for using the same cross-validation folds across all keypoint sets. Following significant Friedman results, we performed pairwise comparisons using the Nemenyi post-hoc test [40].

Our results showed minor performance differences across behaviors and keypoint sets (Figure 1). These minor F1 differences varied from (0.02-0.16) depending on behavior. For instance, Left Turn ranged from F1=0.74 (Mouse Resource, 5 keypoints) to F1=0.76 (JABS, 12 keypoints, p=0.4276, NS). Whereas JABS Supported Rearing ranged from F1=0.90 (JABS) and F1=0.74 (Mouse Resource, p=0.0005). Most behavior performance ranges were tight, and all keypoint configurations performed within error margins of each other. While some behaviors reached statistical significance (Friedman test, p<0.05 for 5 of 7 behaviors), the practical magnitude of these differences was small. A researcher choosing any of the literature-based keypoint sets could expect strong classifier performance, regardless of their choice.

To quantify whether adding keypoints improves performance, we calculated the Pearson correlation coefficient between F1 and keypoint number across literature-based sets. We report R^2^ for the linear fit, and more importantly, the slope quantifying the change in F1-score per added keypoint. If adding more keypoints substantially improves performance, we would expect a large positive slope. For most behaviors, this analysis showed high R^2^ indicating good linear fit (0.50-0.99), but negligible slopes (0.0032-0.0214, Figure S1a). This means that each additional keypoint improves F1 by less than 0.02 on average, which is a negligible practical improvement. This result added another line of evidence that increasing the number of keypoints does not meaningfully improve classifier performance. Furthermore, the effort to add more keypoints far exceeds any performance boost that adding each keypoint gives.

Increasing the number of keypoints is just one dimension of the question; the computed feature set derived from the keypoints may be what actually affects classifier performance. Importantly, the relationship between number of keypoints and number of features is not always linear (Table 1). Features are computed from keypoint combinations (e.g. pairwise distances, angles, etc), meaning that even a small keypoint set can generate a large feature matrix. We therefore repeated the correlation analysis using feature count rather than keypoint number. We found similar results (Figure S2a). Though R^2^ values (0.611-0.995) indicated strong linear relationships, the slopes were extremely low (*≤* 0.0001, Table S1). Neither keypoint count nor feature count meaningfully predicts classifier performance, which again challenges the assumption that more keypoints yields better behavior classifiers.

While most behaviors showed minimal sensitivity to keypoint selection, we examined two behaviors more closely: Scratch (F1 range: 0.77-0.91) and Sturman Supported Rearing (F1 range: 0.81-0.86). For Scratch behavior, differences in performance were driven between the sets that lacked rear paws (MoSeq, F1=0.80; Mouse Resource, F1=0.75), and those that included them (JABS, F1=0.90; MARS, F1=0.89, Table S3). Since Scratch is defined by the process of the mouse using its rear feet to scratch a body part ([5]), the inclusion of these rear paws appeared relevant. On the other hand, even the sets without rear paws performed reasonably well (F1*≥*0.75).

For Sturman Supported Rearing, the differences arose between the full keypoint set (JABS, F1=0.86) and other literature-based sets (MARS, F1=0.83; MoSeq, F1=0.81; Mouse Resource, F1=0.81). Rearing is characterized by the mouse lifting its front paws off of the ground, with supported rearing further defined by rearing against a wall. Rearing is a challenging behavior to quantify from top-down camera views since the mouse occludes the lower half of its body. Notably, lower-performing sets all lack front paw keypoints, which remain visible during rearing, potentially making them particularly salient points for classification.

Across other behaviors (JABS Supported Rearing, JABS Unsupported Rearing, and Sturman Unsupported Rearing), the minimal differences were consistently driven by the most extreme comparison: JABS (12 keypoints) and Mouse Resource (5 keypoints). The majority of pairwise comparisons showed no significant differences, supporting the conclusion that most keypoint configurations yield comparable performance. Even the statistically significant differences only have a small effect size, raising the question: what amount of F1 difference constitutes a meaningful reduction for a useful classifier?

### 2.2 Behavior Classification Performance is Robust to Variation in Systematically Ablated Keypoint Sets

Given that literature-based keypoint sets showed a minimal relationship between keypoint number and performance, we followed up on this idea by testing even smaller keypoint sets. We systematically removed keypoints (ablate) down to an extreme 2-keypoint set (Nose and Base Tail). We hypothesized that at some point in our ablations, we would have a drop off in performance. We predicted that complex behaviors such as rearing would be more sensitive to keypoint selection, while simpler turning behaviors would be more resistant as they rely primarily on angular changes in the body as a whole.

The keypoint sets are described in Table 1. We designed realistic sets by bilaterally removing body parts (paws, ears, tail, etc). We reasoned these sets reflected choices experimentalists might make. Our one exception was No Nose or Tail, which tests the impact of the commonly labeled nose keypoint. We evaluated these ablations using the same experimental framework as the literature-based keypoint sets.

Our ablation analysis showed minimal performance differences across most keypoint configurations (Figure 1b). Mid-sized keypoint sets (6-10 keypoints) performed nearly identically to the full 12-keypoint set, with differences driven by informative keypoints, rather than quantity. Turning behaviors showed minimal F1 variation: Left Turn varied by only 0.04 (F1: 0.72-0.76) and Right Turn by 0.06 (F1: 0.81-0.87). Even for the more complex rearing behaviors, the top-performing sets showed tight clustering. Specifically, for JABS Supported Rearing, the three best performers (No Tail, No Nose or Tail, and Nose and Paws) had F1 scores between 0.89-0.91 despite the variation in keypoint count from 6-10 keypoints. Notably, Nose and Paws (6 keypoints) outperformed No Paws (8 keypoints), showing that the informativeness of the keypoint selection can be more important than quantity.

**Figure 1:**
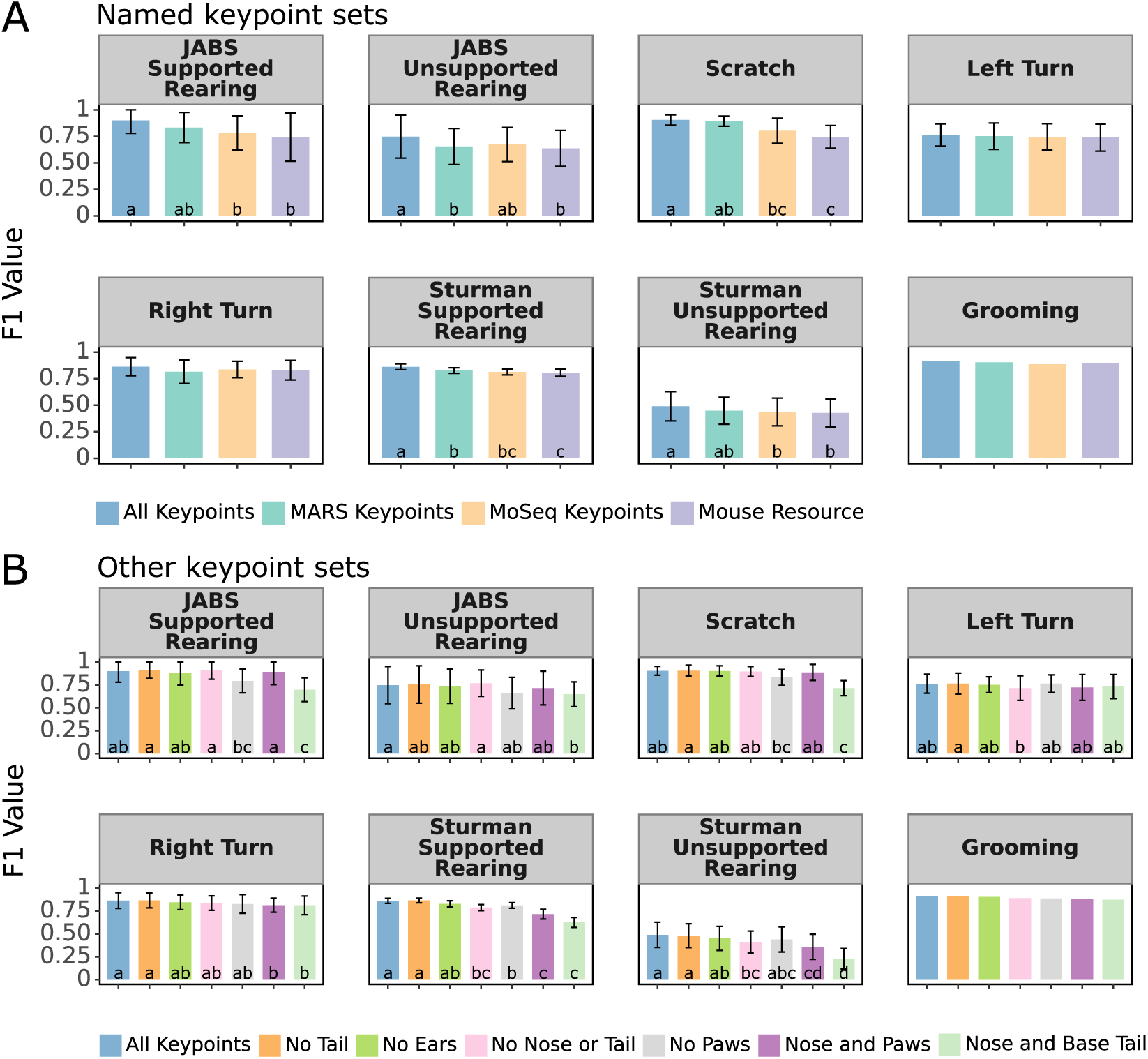
Behavior classifiers are largely invariant to keypoint selection. (A) F1 scores for eight behaviors across four literature-based keypoint sets (JABS, MARS, MoSeq, and Mouse Resource). (B) F1 scores across systematic ablations to test impact of removing specific body parts. Error bars indicate standard deviation across cross-validation folds. Grooming behavior lacks error bars as it uses the fixed held-out validation set from [38]. Statistical comparisons performed using Friedman test with Nemenyi post-hoc comparisons. Letters indicate compact letter display ([42]). Sets sharing a letter are not significantly different (p*≥* 0.05). For example, sets labeled "a" and "ab" do not differ significantly, but sets labeled "a" and "b" do.

Examining behaviors individually shows these patterns more clearly. For Sturman Supported Rearing (excluding the 2-keypoint set) the performance ranged from F1=0.71 (Nose and Paws) to F1=0.87 (No Tail), with All Keypoints (F1=0.86) performing equivalently to No Tail. Sturman Unsupported Rearing had less variation, albeit with lower performance across the board (full keypoint set F1<0.5). However, even in this case (excluding the 2-keypoint set), variation remained limited (F1 range: 0.36-0.49). Nose and Base Tail had the lowest performance, with an F1 of 0.23.

As we expected, we found that the major differences in performance were driven by the extreme 2-keypoint set (Nose and Base Tail). However, even this minimal representation reached reasonable performance: F1=0.70 for JABS Supported Rearing, F1=0.65 for JABS Unsupported Rearing, and F1=0.72 for Scratch. For simple turning behaviors, this extreme set performed within the error margins of the more complex sets (F1=0.72-0.81). The fact that a 2-keypoint set, which is an 83% reduction in keypoints from the 12-keypoint set, maintained performance drops of only 0.03-0.26 compared to the full keypoint set across behaviors shows how efficiently classifiers can extract behavioral signal from limited pose data.

Scratch behavior is an interesting case, with Nose and Base Tail (F1=0.72) having differences from all sets except No Paws (F1=0.83). No Paws also only differs from the top performer No Tail (F1=0.91), but does not differ from any other set. Since scratching is defined by the rear foot movements, the exclusion of paws impacted performance marginally. This again indicates that having informative keypoints for the particular behavior can be more impactful on performance than just keypoint number or feature count. Yet, even this behavior-relevant deficit remained small.

To quantify whether adding keypoints improves performance across our ablated keypoint sets, we calculated Pearson correlations between F1 scores and keypoint number. The slopes were negligible: 0.0036-0.0257 with all sets included, and 0.0059-0.0228 excluding the 2-keypoint set (Figure S1b). This means that each additional keypoint improves F1 by less than 0.02 on average, confirming that adding keypoints does not yield meaningful performance gains. Similarly, correlations between F1 and feature count showed extremely low slopes (m *≤* 0.0001 in all cases, Figure S2b, Table S1). These results confirm that adding keypoints does not result in meaningful performance improvements across the practical range of keypoint sets.

Collectively, these findings challenge the assumption that "more is better" for keypoint annotation. We demonstrate that behavior classifiers are highly efficient at extracting signal from limited pose data. Across the practical keypoint sets (6-12 keypoints), performance is stable. Even at the extreme 2-keypoint set, classifiers surprisingly maintain reasonable performance. The correlation slopes show that adding keypoints provides negligible benefits. Practically, this implies that researchers can prioritize flexible, lower-cost annotation strategies—focusing on the quantity of training frames rather than the density of anatomical keypoints—without sacrificing the validity of their behavioral readouts.

### 2.3 Temporal Feature Effect on Behavior Classification

Having established that keypoint selection has minimal impact on classifier performance, we next examined how feature engineering (specifically how features are computed from the intermediate representation) creates the biggest performance gains. Up to this point, all analyses used a combined temporal feature set (JABS + FFT, selected based on our results in this section) that aggregates spatial information across time windows. However, we have not yet examined whether this temporal information actually improves performance, or compared different approaches to incorporating temporal context.

Since behaviors are quantifying action sequences, the surrounding context around each frame of video should be explored. These behaviors occur over a sequence of frames, rather than in an instantaneous pose. Scratch involves rhythmic limb movements; rearing requires sustained stance. Therefore, we hypothesized that aggregating information across time windows would capture these dynamics and improve classifier performance beyond instantaneous spatial features alone. To test this systematically, we compared classifier performance using four feature computation approaches: base spatial features and three temporal window feature sets from the literature.

We evaluated all keypoint methods using the full 12-keypoint set to isolate the effect of window features from keypoint selection: Base features [5] compute spatial relationships at each frame independently. These include pairwise distances between keypoints, angles formed by keypoint triplets, instantaneous velocities, and velocity directions. For example, base features can provide the nose-to-tail-base distance at frame *t*. The base features would capture instantaneous pose geometry and movement but contain no information about how these measurements evolve over time. They can detect if a mouse’s rear paw is raised at a given moment, but cannot distinguish whether this is part of a rhythmic scratching motion or a single isolated movement. The classifier would need to learn the temporal information independently.

JABS temporal features [5] aggregate base features using statistical summaries (mean, standard deviation, min, max, etc) across a temporal window of surrounding frames Table S5. Rather than solely providing a single instantaneous measurement to the classifier, JABS condenses this temporal information into summary statistics. For example, instead of just the nose-to-tail-base distance at frame *t*, JABS computes over frames *t-16* to *t+16* the mean distance, the standard deviation, the maximum, etc. This approach follows the windowed feature aggregation framework introduced by Kabra et al. [30], where summary statistics transform temporal sequences into compact temporal descriptors.

JAABA temporal features [30] first compute per-frame transformations, then aggregates these across the temporal window (Table S5). In addition to the basic statistical summaries applied to raw features, JAABA also computes frame-to-frame differences, diff neighbor features, zscore neighbors, hist, harmonic features, etc. Note that while the original JAABA paper [30] used segmentation-based intermediate representations, we apply only JAABA’s temporal aggregation methods to our base features.

FFT (fast fourier transform) temporal features [31] transform feature trajectories into frequency components across time windows (Table S5). This approach calculates power spectral density, skewness, kurtosis, FFT band features, etc. For example, if a mouse’s rear paw is oscillating at 6 Hz during scratching, FFT captures this as high power in the 5-8 Hz band, which encodes the rhythmic patterns into the features instead of requiring the classifier to learn it from the raw position sequences. Therefore, this method can be particularly useful for behaviors with periodic characteristics.

These approaches represent different philosophies for incorporating temporal context. Base features provide instantaneous snapshots, JABS computes statistical summaries across time, JAABA emphasizes rates of change, and FFT shows rhythmic periodicity. All these temporal methods collapse the information from multiple frames into aggregated features, but differ in what they aggregate and how.

In addition to keypoint-based features, we evaluated whether these window features could be applied to segmentation-based feature set. Instead of tracking individual body parts, segmentation represents the animal as a binary mask (every pixel of the mouse) at each frame. From these masks, we extract spatial features analogous to our keypoint-based features (Table S5), then apply the same window features (JABS, JAABA, FFT). Segmentation-based approaches offer a potential advantage in annotation efficiency, particularly from recently published methods like SAM2 [33], which can segment an entire video from a single prompt. However, the segmentation-based methods provide coarser spatial information than keypoints. We included segmentation in this analysis to test whether adding temporal window features could compensate for the reduced detail in body features. The full comparison of keypoint versus segmentation representations is presented in Section 2.4.

Temporal features consistently improved classification performance over base features across behaviors (Figure 2). For keypoint-based features, temporal sets improved F1 scores by 7-13% on average: JABS improved by 7.1% ±3.7%, JAABA by 8.0% ±3.8%, and FFT by 8.7% ±6.1%. Surprisingly, segmentation-based representations (described in detail in Section 2.4) showed even larger improvements from temporal features: JABS improved by 16.3% ±6.0%, JAABA by 14.4% ±6.9%, and FFT by 17.6% ±10.0% (Table S4). With segmentation’s greater benefits from the inclusion of window features, it begins to close the performance gap between the keypoint and segmentation-based classifiers. This introduces the idea that keypoints may not even be necessary at all. This also made sense since even the extremely limited 2-keypoint set performed within the range of the other keypoint sets.

**Figure 2:**
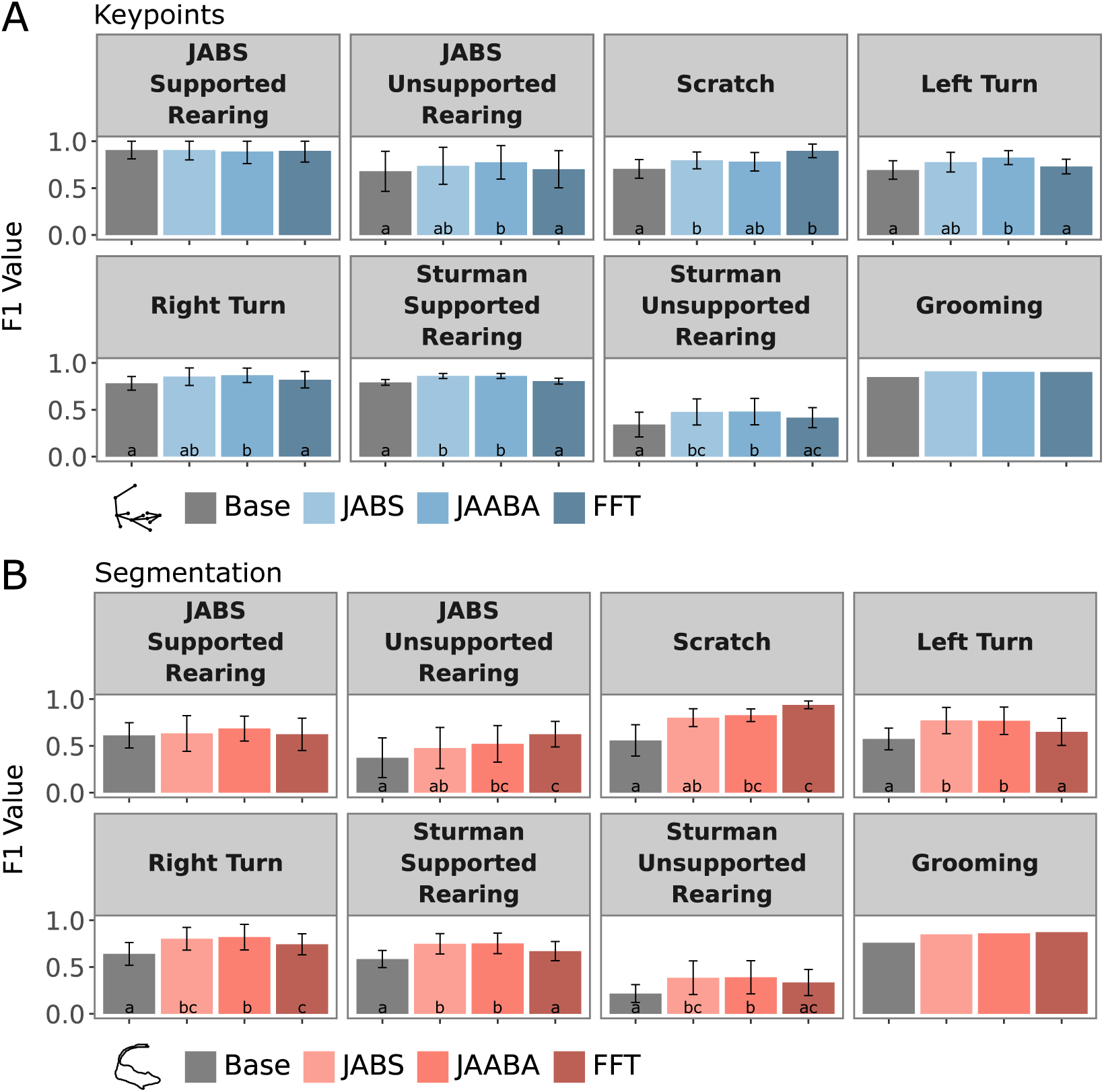
Behavior classifier performance improves with inclusion of window features. (A) Bars indicate F1 scores for keypoint-based features across eight behavior benchmarks, comparing different window feature sets (Base, JABS, JAABA, FFT). (B) Bars indicate F1 scores for segmentation-based features across eight behavior benchmarks, comparing different window feature sets (Base, JABS, JAABA, FFT). Error bars indicate standard deviation across cross-validation folds. Grooming behavior lacks error bars as it uses the fixed held-out validation set from [38]. Statistical comparisons performed using Friedman test with Nemenyi post-hoc comparisons. Letters indicate compact letter display ([42]). Sets sharing a letter are not significantly different (p*≥* 0.05). For example, sets labeled "a" and "ab" do not differ significantly, but sets labeled "a" and "b" do..

The amount of improvement from temporal features varied across behaviors. For Scratch, temporal features improved keypoint F1 from 0.71 (base) to 0.90 (FFT), and segmentation F1 from 0.56 (base) to 0.94 (FFT), with segmentation actually outperforming keypoints. Similarly, JABS Unsupported Rearing showed large improvements from 0.68 (base) to 0.77 (JAABA) for keypoints, and 0.37 (base) to 0.62 (FFT) for segmentation. The larger improvements in the complex behaviors showed the value of the temporal context in analysis of action sequences.

In contrast, JABS Supported Rearing showed no significant differences. Interestingly, this behavior already achieved high performance with solely the keypoint base features (F1=0.91). This suggests that adding temporal features may provide diminishing returns when the base features are already highly discriminative.

Across the three temporal feature sets (JABS, JAABA, FFT), the best performer varied by behavior with no consistent winner. As a combined (JABS + FFT) feature set performed the most consistently across behaviors (Table S4), we used this set for all analyses in other sections of this manuscript. This choice prioritizes robustness across diverse behaviors over optimizing for any single behavior. We find that testing temporal features shows obvious improvements to classifiers, and researchers should continue to engineer more time-based features to improve classifiers.

This agreement of window feature sets improving performance for both keypoint and segmentation suggests that the temporal context is critical to behavior classification, and can be even more important than the specific body parts themselves. Researchers can actually achieve performance increases, particularly for segmentation-based feature sets where there is lot of savings in terms of time-cost, by incorporating temporal features into their feature design. The promising performance of segmentation with temporal features raises the next question: are keypoints even necessary? Segmentation requires substantially less annotation effort (potentially a single prompt per video, without a specialized model trained), yet with appropriate temporal features reaches comparable performance. We explore this alternative approach systematically in the next section.

### 2.4 Segmentation is a Viable Alternative to Keypoints

Developing generalizable, robust keypoint models requires high time-cost and effort for data annotation. With recent advances in segmentation foundation models such as SAM2 [33], obtaining high-quality segmentation masks has become a much cheaper alternative, requiring only a single prompt per video. We therefore asked if keypoint-based features are even necessary, or if there is enough information in carefully crafted segmentation-based features to create comparable, high-quality behavior classifiers.

To address this question, we compared segmentation-based features against keypoint-based features using the same classifier training and evaluation protocols as previous sections. Specifically, we used the JABS and FFT combined temporal feature set for all representations to ensure that the performance differences are reflected by the body representation, rather than the temporal feature set choice. For keypoint-based features, we tested all sets from the previous sections to establish performance range. We compared segmentation-based features against this performance range using Friedman tests followed by Nemenyi post-hoc tests.

We found that segmentation-based features are a viable alternative to keypoint-based features across all tested behaviors (Figure 3a). When compared against the four literature-based keypoint sets (Table 1), segmentation performance was statistically equivalent to at least one keypoint configuration for every behavior (Friedman test with Nemenyi post-hoc, p *≥* 0.05). This shows that segmentation is consistently within the performance range of keypoint subset selection.

**Figure 3:**
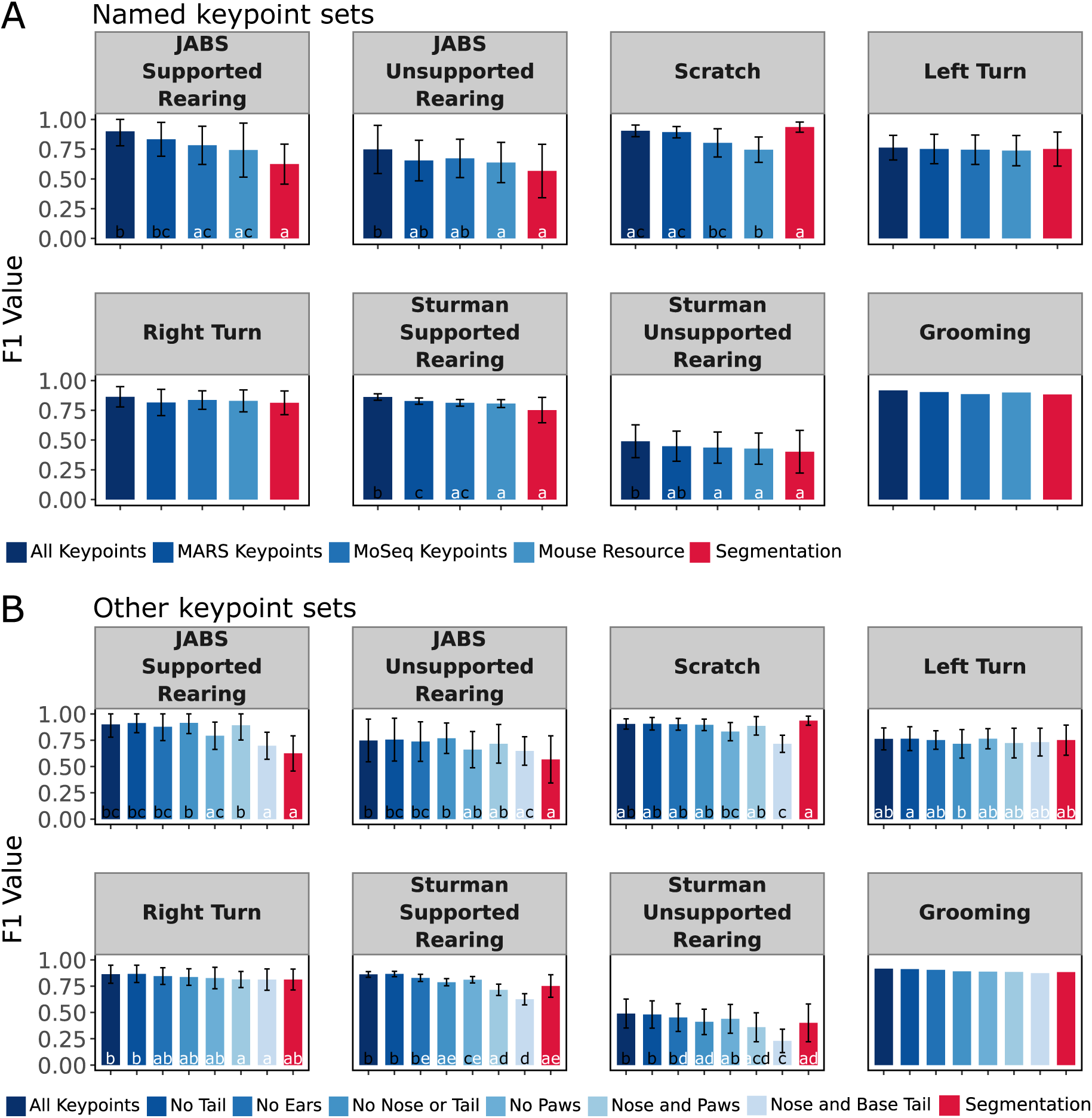
Segmentation-based classifiers are comparable with keypoint-based classifiers. (A) F1 scores for eight behaviors across four literature-based keypoint sets (JABS, MARS, MoSeq, and Mouse Resource) compared to segmentation-based classifier (red). (B) F1 scores across systematic ablations to compare sparser keypoint-based feature sets against segmentation-based classifier. Keypoint sets are in varying shades of blue, segmentation is highlighted in red. Error bars indicate standard deviation across cross-validation folds. Grooming behavior lacks error bars as it uses the fixed held-out validation set from [38]. Statistical comparisons performed using Friedman test with Nemenyi post-hoc comparisons. Letters indicate compact letter display ([42]). Sets sharing a letter are not significantly different (p*≥* 0.05). For example, sets labeled "a" and "ab" do not differ significantly, but sets labeled "a" and "b" do.

The relationship between segmentation and keypoint performance varied by behavior and behavior complexity. For the simple turning behaviors, segmentation performed equivalently to all keypoint sets: Left Turn (Friedman p=0.60) and Right Turn (Friedman p=0.20) showed no statistical differences between any groups, including segmentation.

For the more complex rearing behaviors, segmentation typically grouped with the mid-sized keypoint sets, rather than the full 12-keypoint set (Figure 3a). For both Unsupported Rearing behaviors (JABS and Sturman), segmentation was statistically equivalent to MoSeq and Mouse Resource sets (5-6 keypoints). For both Supported Rearing behaviors, segmentation additionally grouped with MARS (10 keypoints), differing significantly only from the full All Keypoints (JABS) set. Segmentation performs comparably to the reduced keypoint sets while falling slightly short of the densest keypoint set for these complex behaviors.

We found that segmentation matches or exceed keypoint performance for specific behaviors. Strikingly, segmentation had the highest performance for Scratch (F1=0.94), forming a top-performing statistical group alongside JABS (F1=0.90) and MARS (F1=0.89), while significantly outperforming the two sparsest keypoint sets: Moseq (F1=0.80) and Mouse Resources (F1=0.75). This result shows that for certain behaviors, segmentation-based features can capture patterns as effectively, or even better than keypoint configurations.

To further understand the range of the performance of segmentation, we compared it against the sparser keypoint sets with specific keypoints bilaterally removed (Table 1). Segmentation remained statistically equivalent to multiple keypoint sets for all behaviors (Figure 3b). For JABS Supported and Unsupported Rearing, segmentation grouped with all sets that excluded paws, showing its comparable performance to minimal keypoint sets. Again, for Scratch, segmentation outperformed all other sets (F1=0.94), grouping with the higher performing ablations and only significantly differing from No Paws (F1=0.83) and Nose and Base Tail (F1=0.72).

For Sturman Rearing behaviors, segmentation grouped with mid-range ablations, consistently falling between the best and the worst performers. Again, for the turning behaviors (Right Turn and Left Turn), segmentation showed no statistical differences from any keypoint-based set.

Notably, though segmentation ranked the lowest for JABS Supported and Unsupported Rearing, it remained statistically equivalent to several keypoint sets, showing that even lower mean performances does not necessarily indicate meaningful practical differences.

These results show that while carefully selected keypoint sets may provide marginally better results for certain behaviors, segmentation represents a viable and often equivalent alternative. The consistent segmentation performance across the keypoint selection range makes it particularly valuable where a single feature set must generalize across multiple behaviors, or the optimal keypoint set is unknown at the time of model training. Segmentation is also valuable to consider as the annotation cost is now substantially lower than keypoints with methods such as SAM2 [33] that only requires a single prompt per video.

### 2.5 Classifier Performance Scales with Annotation Dataset Size, Making Annotation Quantity More Valuable Than Keypoint Refinement

The grooming dataset is large, with over 2 million annotated frames, making it a good choice for inspecting how classifier performance scales with training data across various feature sets. This analysis has practical implications, since many behavior labs cannot afford to invest annotation time required to label datasets of this size and have to decide whether to prioritize keypoint model refinement or adding more behavioral annotations.

We subsampled the grooming training set at multiple scales with multiple replicates per size. We evaluated performance for temporal features, named keypoint sets, and the other keypoint ablations. We then assessed how these different representation choices interact with training data quantity.

Across all comparisons, increasing annotation dataset size improved the classifier performance regardless of feature set (Figure 4a). This confirms the unreasonable effectiveness of training data [43, 44]. Critically, this improvement occurs for all keypoint sets, temporal feature types, and even segmentation-based representations. The choice of representation can minimally affect absolute performance levels, but all sets benefit substantially from additional behavior annotations.

**Figure 4:**
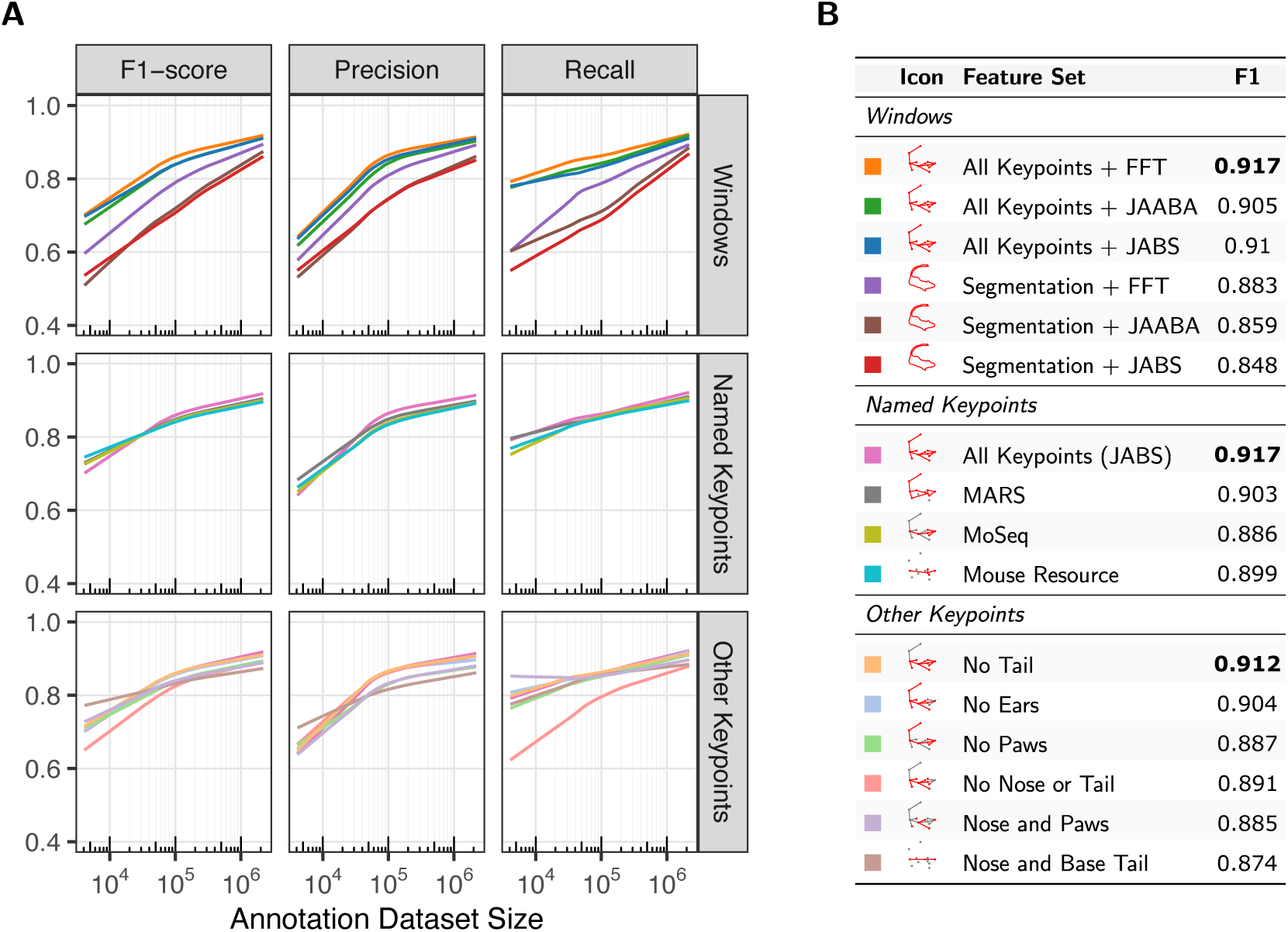
All feature sets detail performance improvement as grooming behavior annotation dataset size increases. (A) Grooming behavior performance scaling curves comparing window-based feature groups, literature-based, and other systemic ablation sets across increasing annotation dataset sizes. (B) Summary table of F1 scores on the full grooming dataset. Bold indicates best performance for each category.

First, we look at how temporal feature sets are impacted by training dataset size. Interestingly we found that despite providing over 2x more features than JABS, the FFT feature set was not adversely affected by overfitting at small training dataset sizes (Figure 4a, top row). All three temporal feature sets (JABS, JAABA, FFT) converged to similar performance at the full dataset size of 2.18 million frames, with FFT achieving F1=0.91 for All Keypoints and F1=0.88 for Segmentation (Figure 4b). When comparing keypoint versus segmentation, we find an important result (Figure 4a, top row) At small dataset sizes, keypoints consistently outperform segmentation. However, segmentation shows a steeper learning curve and closes the performance gap as the behavioral annotations increase. This scaling behavior has important practical conclusions since segmentation only requires a single prompt per video using methods like SAM2 [33], compared to training a specialized keypoint model with all body parts that the user wants to track being included in the time cost. This means that the behavioral annotation budget can be used to determine the final performance. Since we see that performance starts to level off around 10^5^ annotated frames for both approaches, investing annotation effort in behavior labels rather than refining keypoint models shows a more reliable path to strong performance.

We next examined how keypoint selection itself interacts with dataset size. At larger dataset sizes (>10^5^ frames), keypoint set selection has minimal impact on performance (Figure 4a, middle and bottom rows), consistent with our previous findings. However, as training dataset size decreases, both the best performing keypoint set and the performance variation across sets become increasingly variable (Supplemental Figure S3, Supplemental Figure S4). At the smaller dataset sizes (10^4^), keypoint sets varied in performance ranking. This creates a risk of choosing the wrong keypoint set, and determining the optimal set beforehand is challenging without substantial behavioral annotation to evaluate performance.

Exploring keypoint-based representations at even smaller subsets reveals that a good performing grooming behavior classifier is attainable with only 2 keypoints (Nose and Base Tail). Nose and Base Tail had an F1 of 0.87 on the full dataset, competitive with the other more complex keypoint sets (Figure 4a, bottom row; Figure 4b). Interestingly, this minimal set actually had the highest performance at the smallest dataset size (10^4^ frames), though this ranking changed as more data was added. This a suggests that a small number of informative keypoints may be sufficient when annotation resources are limited. However, predicting which minimal set will perform best remains difficult without testing.

These results show that adding behavioral annotations provides more predictable and consistent performance gains than optimizing keypoint selection. At small dataset sizes, keypoint choice matters more but is unpredictable, while at larger dataset sizes, the keypoint choice hardly matters at all. In contrast, adding more behavioral annotations consistently improves performance across all representations. For labs with limited annotation resources, prioritizing the quantity of behavioral labels over keypoint model refinement offers a more robust strategy. This is particularly true when using segmentation-based approaches that substantially minimize the pose estimation annotation costs.

## 3 Discussion

The transition from manual scoring to automated behavior quantification promises to transform neuroethology by enabling scalable, objective, and reproducible measurements. Rather than relying on manual scoring which is slow, subjective, and un-scalable, or collapsing all behavior into solely summary measures such as total locomotor activity, automated classifiers can provide reproducible measurements of specific behaviors across large cohorts of animals. This is particularly impactful for high-throughput applications such as therapeutic screening, where behavior is an important facet of determining whether drug candidates or other interventions have the intended effects. Despite this potential, the practical barriers to building and applying these classifiers remain high. The annotation burden required to train pose estimation models and behavior classifiers limits who can participate, and has pushed many laboratories toward oversimplified behavioral measurements that sacrifice specificity and granularity. This bottleneck has been exacerbated by a prevalent, yet largely untested, assumption that denser keypoint sets, i.e. a higher number of keypoints, produces better behavioral classification. This reasoning is intuitive as more anatomical and fine-grained information should, in theory, provide richer features for classification. Our findings challenge this orthodoxy. By systematically benchmarking intermediate representations, we demonstrate that the "more is better" intuition for pose estimation yields diminishing returns. Instead, we show that optimizing the annotation strategy—prioritizing behavioral volume over keypoint density and leveraging lower-cost segmentation representations—offers a more efficient path to high-accuracy quantification.

Our results indicate that classifier performance is remarkably robust to keypoint selection. While the field has trended toward denser keypoint sets (e.g., TopViewMouse-5K (27 points) [45], JABS (12) [5], MARS (9) [10], MPD-OFT (7) [46], MoSeq (6) [14], Mouse Resource (5) [37]), we find that increasing keypoint counts provide only marginal gains once a minimal anatomical structure is captured. We also find that these marginal performance gains vanish as behavioral training dataset size increases. This has direct implications for how laboratories should allocate their annotation resources. Given that keypoint annotation is costly—estimated at 0.90–1.77 seconds per point [4]—our data suggests that this effort is often misallocated. Adding behavioral annotations, by contrast, is cheaper (0.16-0.25 seconds [4]) and provides consistent, predictable improvements, regardless of the representation used. Laboratories with limited annotation budgets may benefit by investing in behavior annotations rather than expanding or refining their keypoint models. This shift in resource allocation provides consistent, predictable improvements in classifier performance regardless of the underlying representation.

Another practical implication of classifier performance being invariant to keypoint selection is that labs do not need to redesign their pose estimation pipeline every time they wish to study a new behavior. A keypoint model already in use for one set of behaviors is likely to generalize well to new ones. This means that the upfront investment in a pose model can be made once, then reused as the behavioral questions a lab asks continue to evolve. This lowers the effective cost of behavior discovery and can encourage labs to characterize a richer set of behaviors than they otherwise might attempt.

The dominance of keypoint-based representations in modern behavior classification pipelines emerged with the rapid development of pose estimation tools, and has largely remained unquestioned. The dominance of pose in modern pipelines has often been assumed to be superior to segmentation, likely due to comparisons with older, simpler segmentation descriptors [47, 30]. Our results suggest it is time to revisit that conclusion. We show that when we extract a richer set of features from segmentation masks (e.g., Hu moments [48]) and temporal features, they achieve performance parity with keypoint-based classifiers across majority of the behaviors tested. This finding identifies the bottleneck not as the representation itself (segmentation vs. pose), but as the richness of feature extraction.

This distinction matters because the cost of obtaining segmentation has plummeted. The advent of foundational models like SAM2 [33] have effectively automated the segmentation process, reducing the annotation burden by orders of magnitude compared to training custom pose models. For example, generating predictions for the Sturman dataset [4] via SAM2 required only 20 prompts, when compared to the 1560 keypoint annotations (13 keypoints across 120 frames) used in training the Sturman keypoint model, this is an intermediate representation cost savings of 78*×*. This figure is specific to the Sturman dataset and will vary with dataset size and structure, but it illustrates the order-of-magnitude difference in annotation burden that segmentation can offer. There has also been a lot of progress from groups that have worked to fine-tune SAM-based segmentation approaches to mouse tracking [49, 50, 51]. SAM3 [34] was released recently, which enables automated segmentation of mouse masks from a text-prompt alone. This has not yet been benchmarked in lab mice, but seems extremely promising, only furthering the recommendation that the field lean into segmentation-based representations. Segmentation is also a more objective annotation task, which addresses concerns about inter-annotator variability for keypoint placement reported in [10].

Beyond spatial representations, our work highlights the under-utilized power of temporal feature engineering. Consistent with early insights from JAABA [30], we find that incorporating temporal context (specifically via FFT-based window features) consistently improves classifier performance. This result shows the opportunity for further development in this underexplored facet of improving classifier performance without requiring more spatial labels. Even more complex techniques such as recurrent neural networks can be incorporated in the future, while keeping pipelines data-efficient.

Finally, the choice of intermediate representation has profound implications for data sharing and reproducibility. The current landscape of behavioral analysis is fragmented; classifiers are rarely portable between labs due to the incompatibility of specific keypoint definitions. While [11] argue that SHAP values allow classifiers to be compared, the classifiers themselves still cannot be easily transferred between groups. This can be attributed to the limited overlap of intermediate representations. Segmentation offers a path toward a "universal" intermediate representation. Unlike keypoints, which are defined by subjective anatomical choices, the animal’s silhouette is an objective physical property. By adopting segmentation, the community could share data (raw videos and behavior labels) that can be processed by any group using off-the-shelf foundation models, bypassing the need to share or retrain specific pose estimators. While efforts to create generalized keypoint foundation models are promising (e.g., SuperAnimal [45]), segmentation currently offers an immediate, low-barrier solution for standardized data exchange.

The field has begun to address the data scarcity problem. Community efforts such as the MABe competition [52] have released large datasets across labs as shared resources, increasing the supply of labeled behavioral data to the community. They are also targeting the task of having classifiers work appropriately across groups. Our findings complement this effort by providing the recommendation to use annotation-efficient intermediate representations and prioritize behavioral labels over refining pose.

Our hope is that this work encourages labs to treat annotation strategy decisions with the same care as experimental design. The choice of intermediate representation, the investment of time into annotation effort between pose or behavior labels, and the selection of temporal features have consequences on classifier performance, annotation cost, and reproducibility of findings. These choices can now be supported by this benchmark as a start to expand to what behavioral neuroscience is able to measure, and at what scale.

## 4 Limitations of the study

Many current tools designed for biology behavior classification do not support segmentation data inputs. As such, both deep learning toolkits and behavioral classifier tools would need to be upgraded to include segmentation information. This would be a substantial infrastructural investment that would need to take place.

While we present experiments using a single animal, the lab animal behavior field has also included social paradigms. More experiments should be conducted for behaviors in a social context. In addition, it is relatively simple to extend keypoint features that describe relationships between two individuals. However, segmentation features are less clearly extendable. Determining whether the segmentation-based features are applicable in social contexts, or whether keypoint representations become more valuable with multiple animals, remains unexplored.

## 5 Resource availability

### 5.1 Lead Contact

Requests for further information and resources should be directed to and will be fulfilled by the lead contact, Vivek Kumar.

### 5.2 Materials availability

This study did not generate new materials.

### 5.3 Data and code availability

The datasets used in analysis for this paper are all publicly available. Pipeline and analysis code related to this manuscript is available on GitHub: https://github.com/KumarLabJax/JABS-feature-experiments.

## 6 Acknowledgments

This work was funded by The Jackson Laboratory Directors Innovation Fund, National Institutes of Health AG078530 (NIA, V.K.), DA041668 and DA048634 (NIDA, V.K.), MH138309 (NIMH, V.K.), and Nathan Shock Centers of Excellence in the Basic Biology of Aging AG38070 (NIA). T32 AG062409 (NIA, J.D.C.). We thank Kumar Lab members for critical feedback. We thank Camille Berger-Liedtka for project coordination.

## 6.1 Author contributions

Contributions are described using the CRediT (Contributor Roles Taxonomy) categories.

- **Conceptualization:** J.D.C., B.G., V.K.
- **Methodology:** J.D.C., B.G.
- **Investigation:** J.D.C., B.G.
- **Formal analysis:** J.D.C., B.G.
- **Visualization:** J.D.C., B.G., V.K.
- **Writing –** original draft: J.D.C.
- **Writing – review & editing:** J.D.C., B.G., V.K.
- **Funding acquisition:** V.K.

## 6.2 Declaration of interests

The authors declare no competing interests.

### Supplementary Material

**Figure S1:**
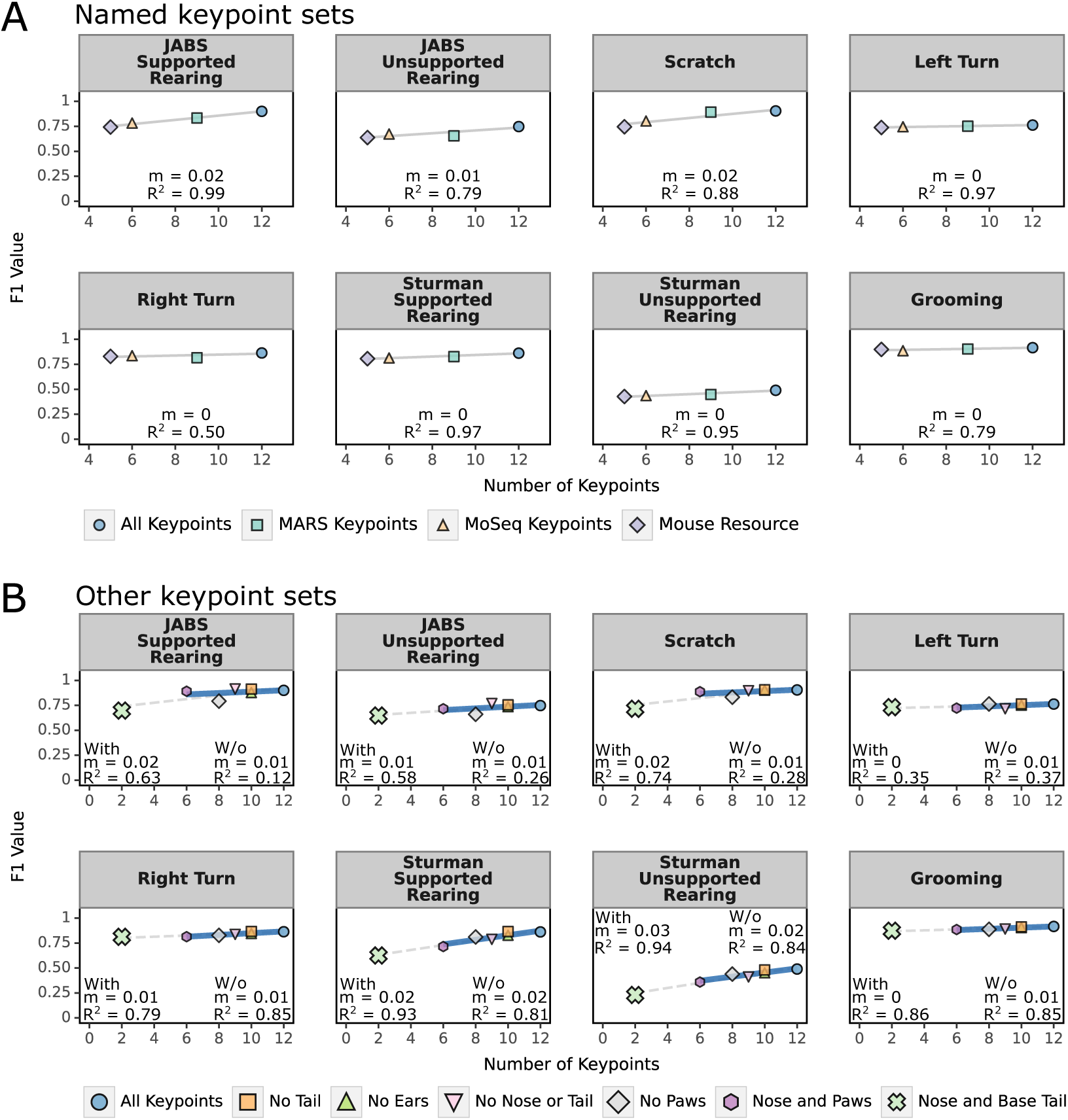
Relationship between keypoint count and classifier F1 performance. (A) F1 scores plotted against number of keypoints for four literature-based keypoint sets across eight behaviors. Each point represents mean F1 score for specific keypoint set. Linear regression lines are shown with slope (*m*) and R^2^ values. (B) F1 scores for keypoint ablations, comparing performance "with" and "w/o" the minimal 2-keypoint set (Nose and Base Tail). Linear regressions are fit separately for the two types of comparisons. y = mx + b, m=slope.

**Figure S2:**
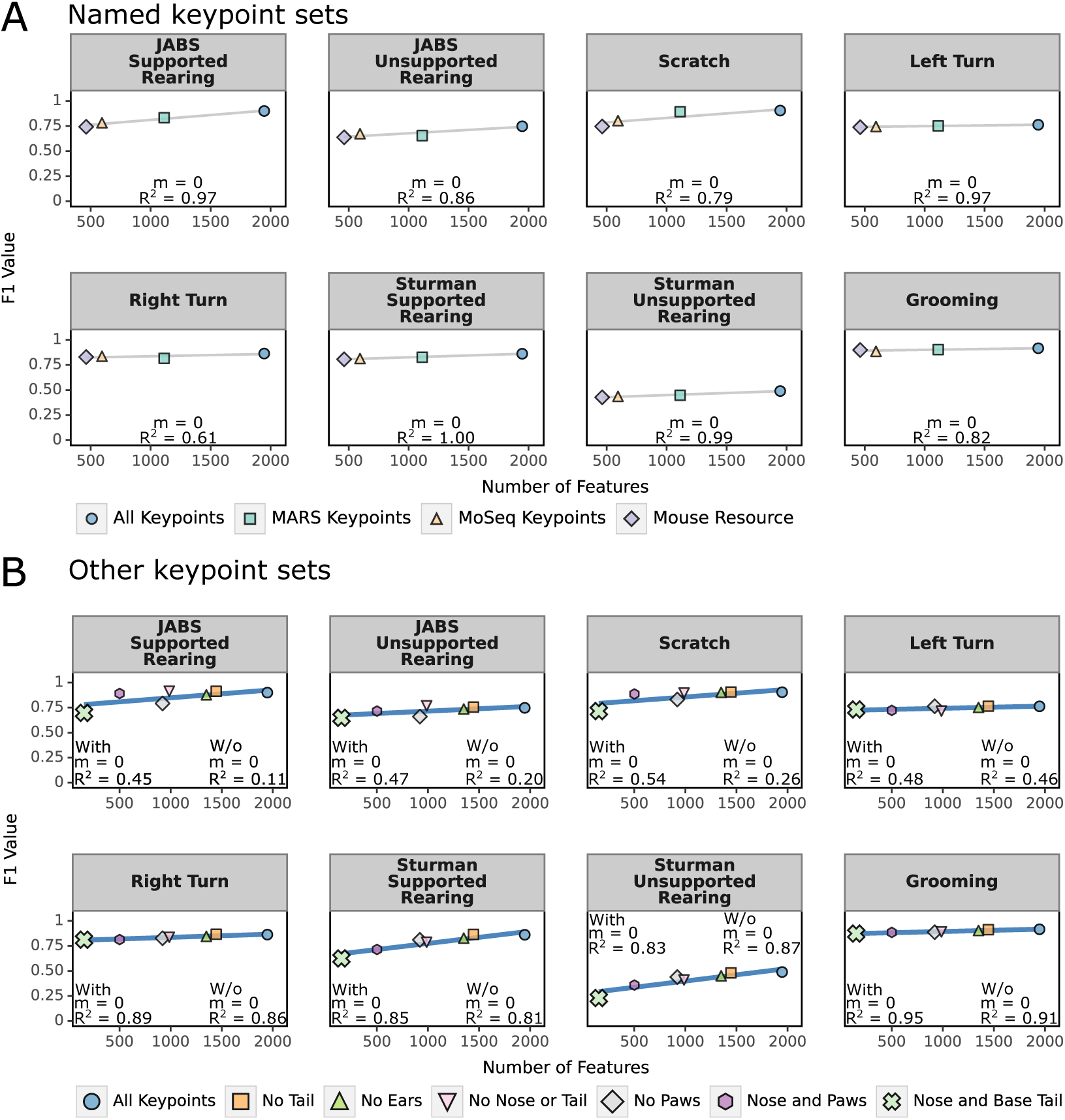
Relationship between feature count and classifier F1 performance. (A) F1 scores plotted against total number of features for four literature-based keypoint sets across eight behaviors. Each point represents mean F1 score for a specific keypoint set after feature extraction (JABS and FFT features combined). Linear regression lines are shown with slope (*m*) and R^2^ values. (B) F1 scores for keypoint ablations, comparing performance "with" and "w/o" the minimal 2-keypoint set (Nose and Base Tail). Points represent different keypoint sets (see legend). Linear regressions are fit separately for the two types of comparisons. y = mx + b, m=slope.

**Table S1:**
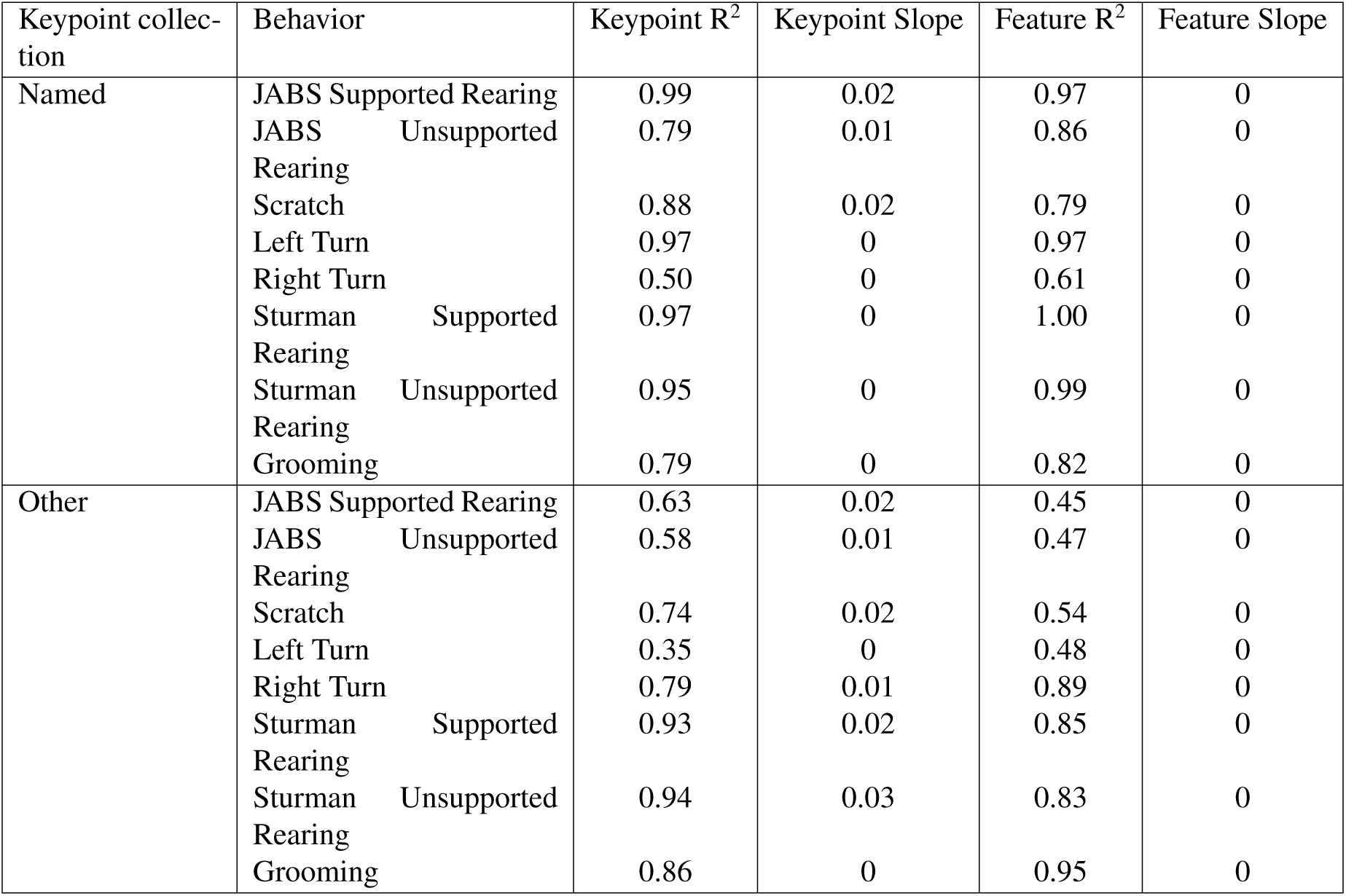
Keypoint and feature Pearson correlation against F1 performance metrics. R^2^ and slopes reported from Figure S1 and Figure S2

#### S0.3 Segmentation Features

In this section, we describe the segmentation features selected for training classifiers. We select only translational invariant descriptors. Image moment description features are calculated using OpenCV [53]. Additionally, we fit an ellipse to the segmentation and use descriptors found in [30]. Additional shape descriptors were calculated based on definitions found in [54]. See Supplemental Table S2 for a full list of included segmentation features.

**Table S2:** Table of segmentation features.

| Feature Name | Feature Description | Group |
| --- | --- | --- |
| hu1 | Hu moment 1 | Image Moment |
| hu2 | Hu moment 2 | Image Moment |
| hu3 | Hu moment 3 | Image Moment |
| hu4 | Hu moment 4 | Image Moment |
| hu5 | Hu moment 5 | Image Moment |
| hu6 | Hu moment 6 | Image Moment |
| hu7 | Hu moment 7 | Image Moment |
| m00 | Area | Image Moment |
| mu20 | Central moment $M_{20}$ | Image Moment |
| mu11 | Central moment $M_{11}$ | Image Moment |
| mu02 | Central moment $M_{02}$ | Image Moment |
| mu30 | Central moment $M_{30}$ | Image Moment |
| mu21 | Central moment $M_{21}$ | Image Moment |
| mu12 | Central moment $M_{12}$ | Image Moment |
| mu03 | Central moment $M_{03}$ | Image Moment |
| nu20 | Scale normalized $\mu_{20}$ | Image Moment |
| nu11 | Scale normalized $\mu_{11}$ | Image Moment |
| nu02 | Scale normalized $\mu_{02}$ | Image Moment |
| nu30 | Scale normalized $\mu_{30}$ | Image Moment |
| nu21 | Scale normalized $\mu_{21}$ | Image Moment |
| nu12 | Scale normalized $\mu_{12}$ | Image Moment |
| nu03 | Scale normalized $\mu_{03}$ | Image Moment |
| cetroid_speed | Magnitude of first derivative of centroid center location | JAABA |
| ellipse_w | Minor axis length of ellipse regression | JAABA |
| ellipse_l | Major axis length of ellipse regression | JAABA |
| perimeter | Length of perimeter | Shape Descriptor |
| elongation | 1 - Length-width ratio of bounding rectangle | Shape Descriptor |
| rectangularity | Area ratio of bounding rectangle and shape | Shape Descriptor |
| convexity | Perimeter ratio of convex hull and shape | Shape Descriptor |
| solidity | Area ratio of convex hull and shape | Shape Descriptor |
| euler_number | Difference of external contours and internal contours | Shape Descriptor |
| hole_area_ratio | Area ratio of shape and its holes | Shape Descriptor |

#### S0.4 Additional Keypoint Subsets

In addition to the keypoint subsets described in [10, 37, 14], we include experiments with an additional 6 keypoints subsets. These subsets can be visualized with the icons in Supp Table 1 where red points are included and gray points are excluded. The "No Ears" subset excludes the 2 ear keypoints. The "No Tail" subset excludes the 2 keypoints on the tail. The "No Paws" subset excludes the 4 paw keypoints. The "No Nose or Tail" subset excludes the 2 tail keypoints and the nose keypoint. The "Paws and Nose" subsets only includes the 4 paw keypoints and the nose keypoint. The most extreme is "Nose and Base Tail", where only the nose and base tail keypoints are included.

#### S0.5 Grooming scaling experiments, extended

The figures in this section are based on Figure 4, with performance metrics including precision, recall, f1, accuracy, and roc auc. These plots include the individual datapoints from the shuffled scaling experiments.

**Figure S3:**
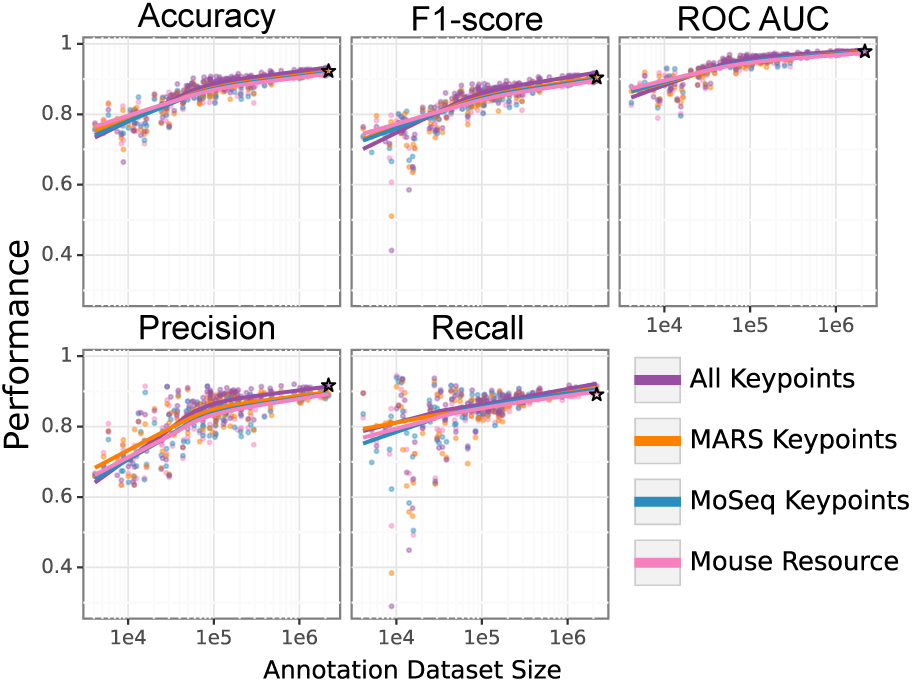
Grooming behavior classifier performance across training dataset sizes and different keypoint subsets. The keypoint sets are the published sets present in Table 1. This is an expanded version from Figure 4. Stars indicate CNN from [38].

**Figure S4:**
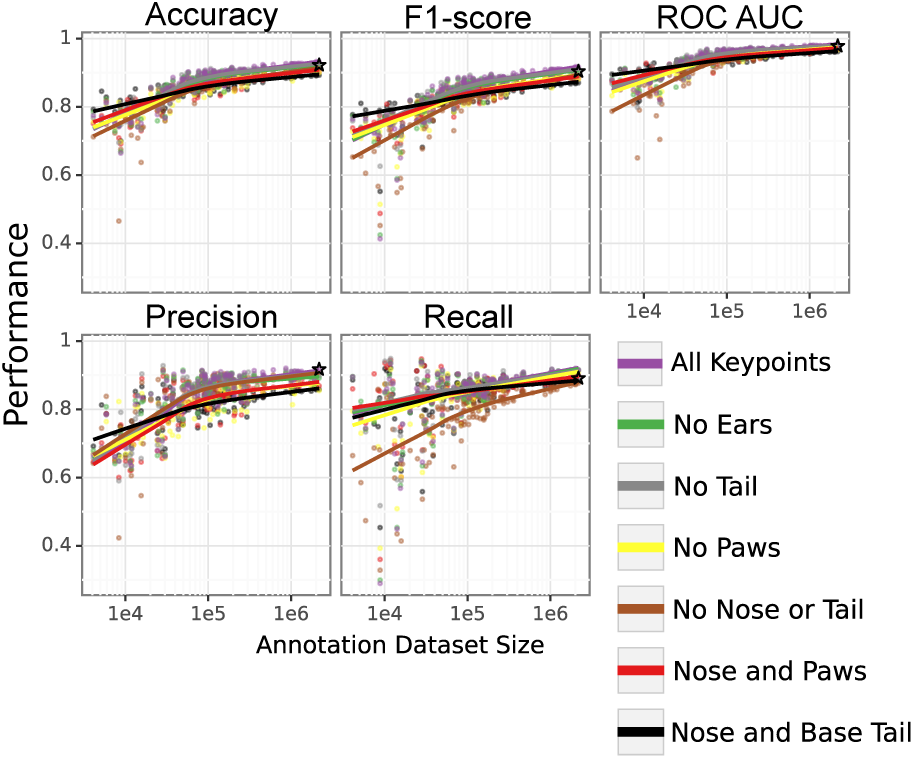
Grooming behavior classifier performance across training dataset sizes and different keypoint subsets. The keypoint sets are the hand-crafted sets present in Table 1. This is an expanded version from Figure 4. Stars indicate CNN from [38].

**Table S3:**
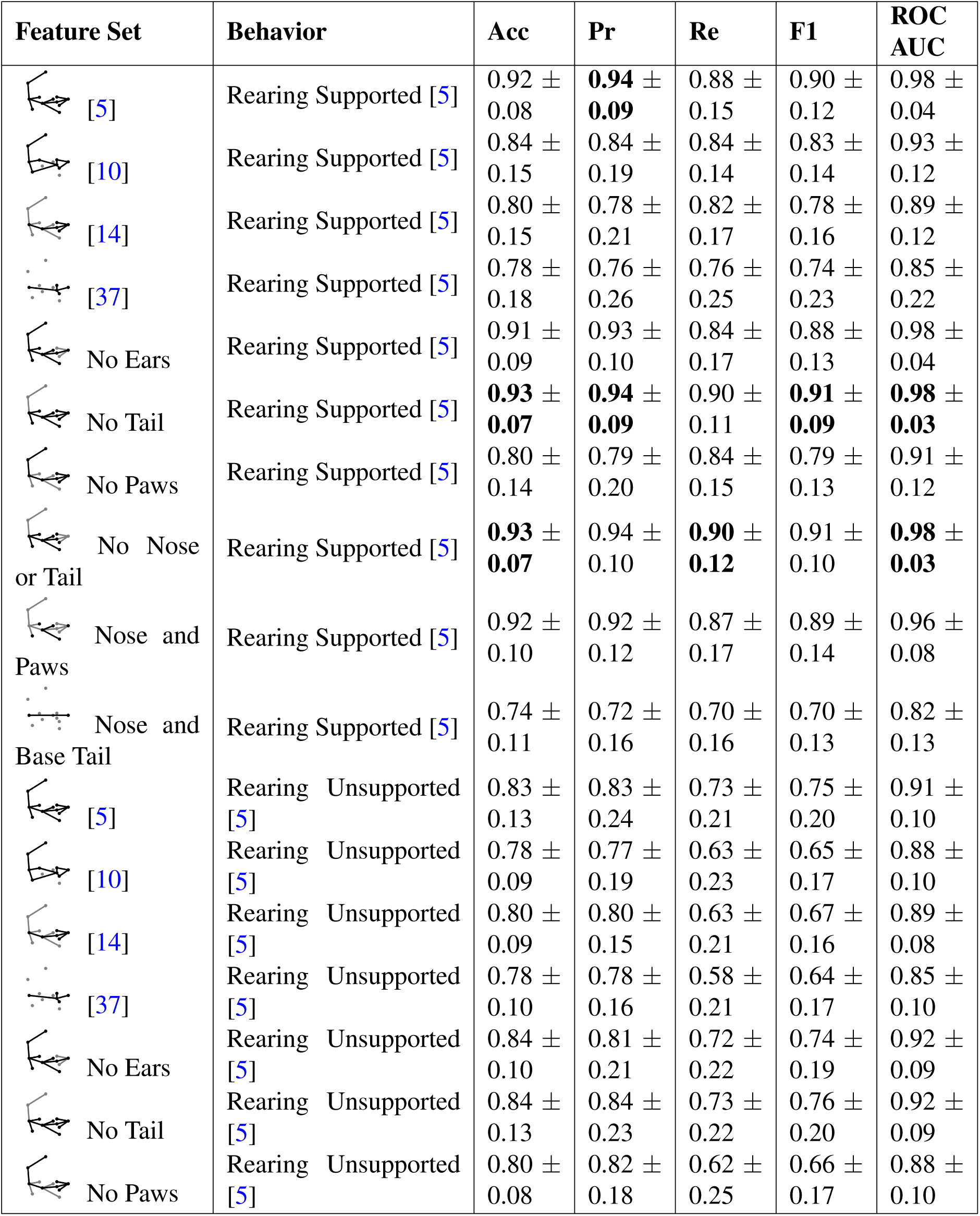
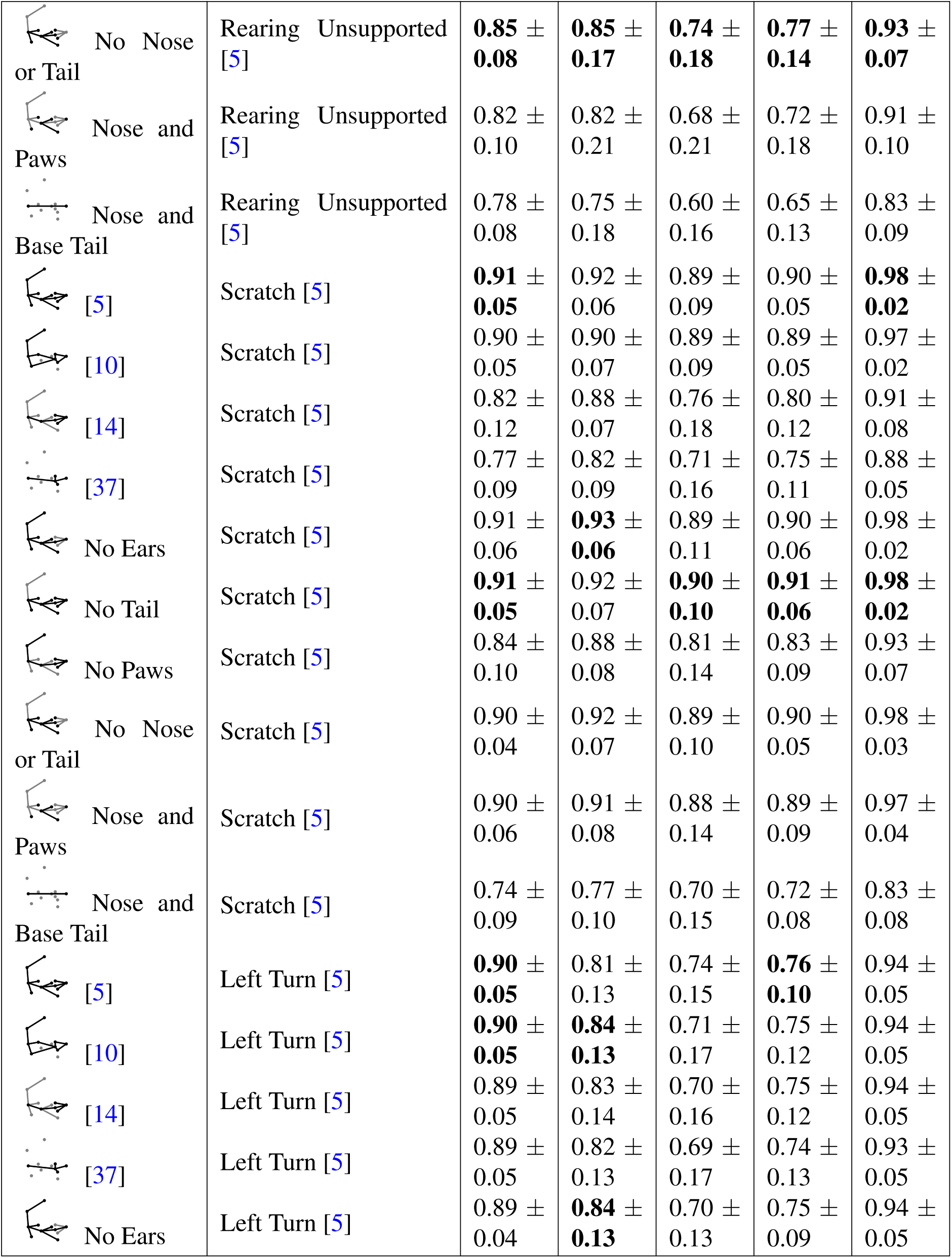
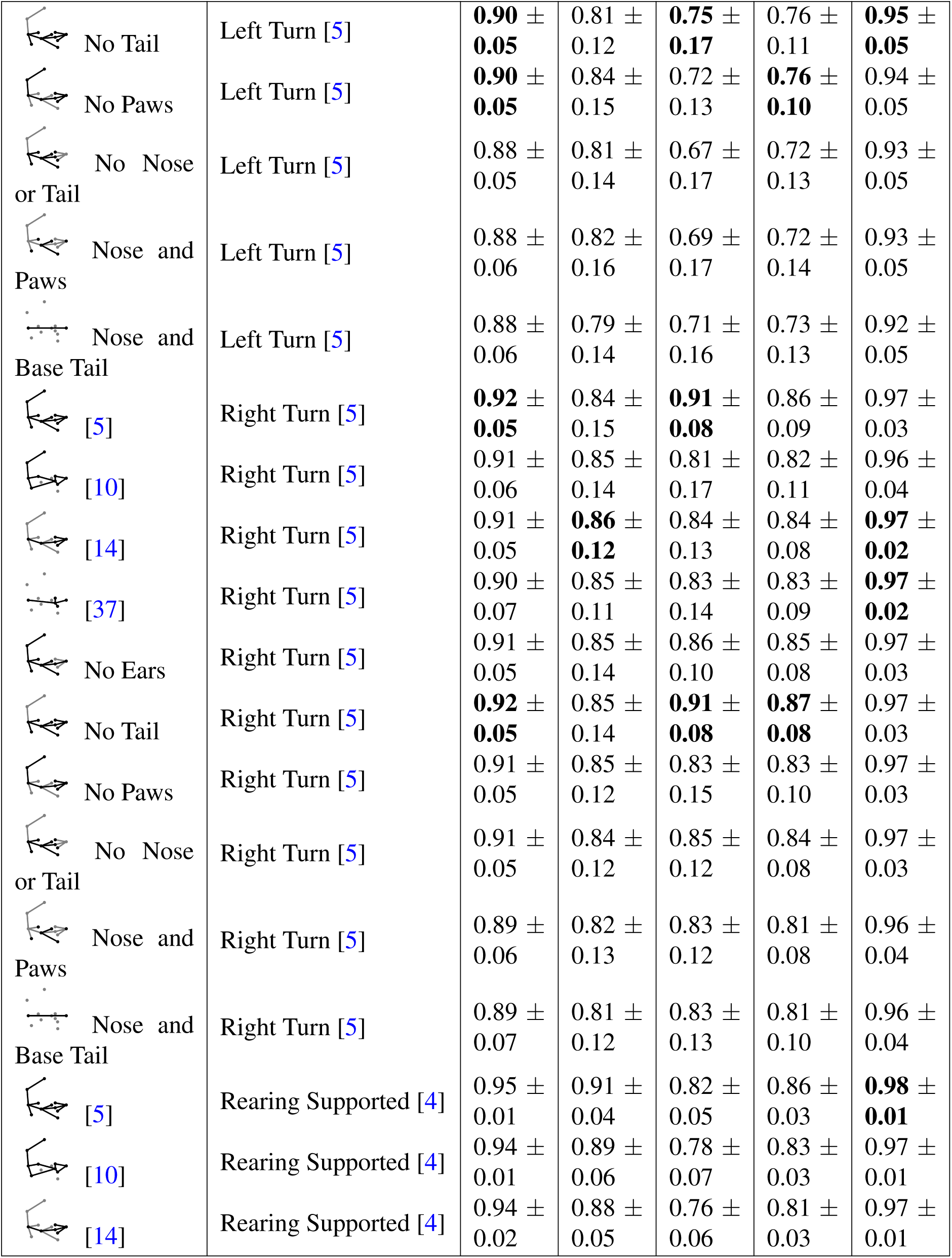
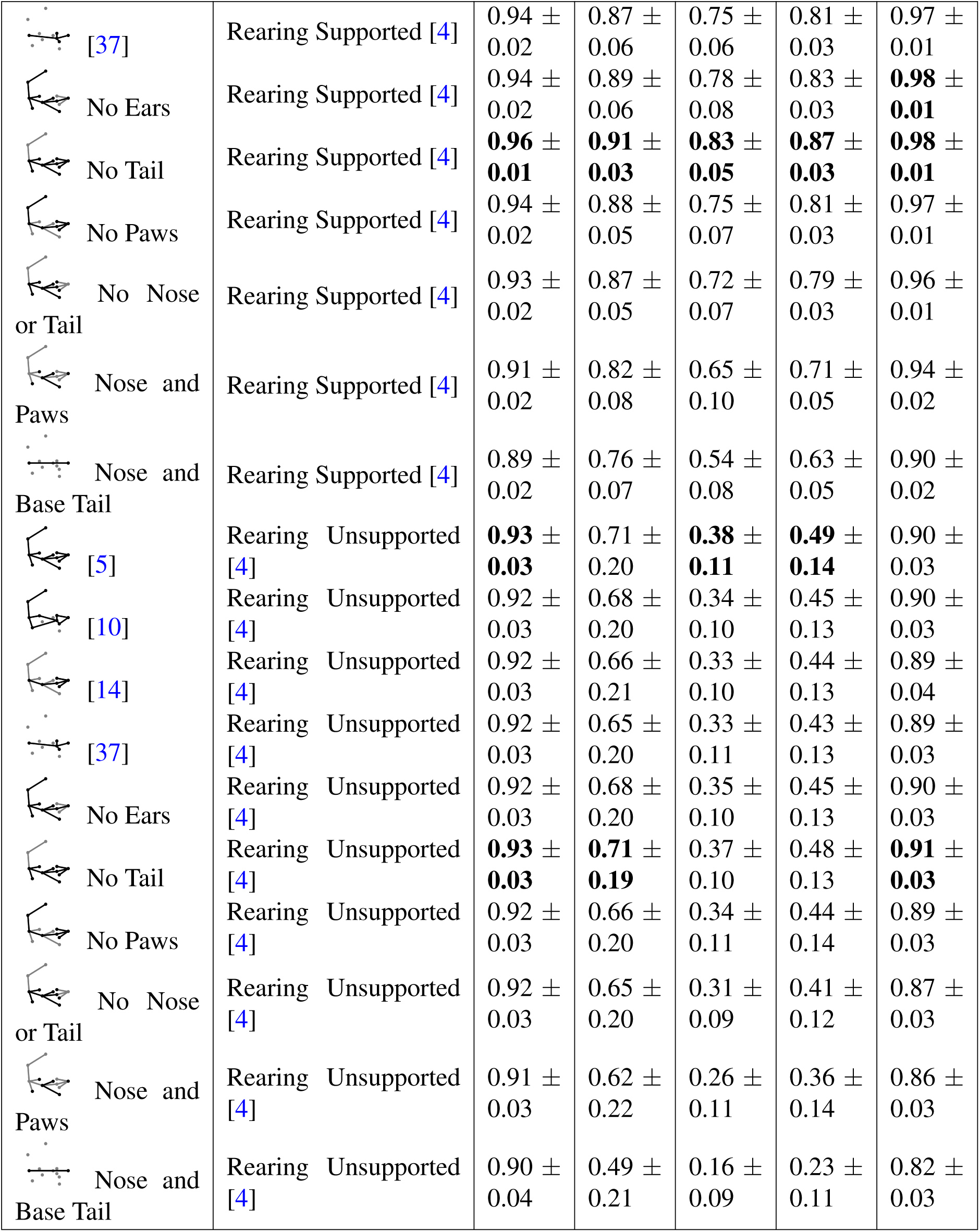
Performance (mean *±* std) of classification tasks across additional behaviors for different keypoint subsets trained with FFT window features. Bold indicates best performance.

| Feature Set | Behavior | Acc | Pr | Re | F1 | ROC AUC |
| --- | --- | --- | --- | --- | --- | --- |
| [5] | Rearing Supported [5] | 0.92 ± 0.08 | **0.94** ± **0.09** | 0.88 ± 0.15 | 0.90 ± 0.12 | 0.98 ± 0.04 |
| [10] | Rearing Supported [5] | 0.84 ± 0.15 | 0.84 ± 0.19 | 0.84 ± 0.14 | 0.83 ± 0.14 | 0.93 ± 0.12 |
| [14] | Rearing Supported [5] | 0.80 ± 0.15 | 0.78 ± 0.21 | 0.82 ± 0.17 | 0.78 ± 0.16 | 0.89 ± 0.12 |
| [37] | Rearing Supported [5] | 0.78 ± 0.18 | 0.76 ± 0.26 | 0.76 ± 0.25 | 0.74 ± 0.23 | 0.85 ± 0.22 |
| No Ears | Rearing Supported [5] | 0.91 ± 0.09 | 0.93 ± 0.10 | 0.84 ± 0.17 | 0.88 ± 0.13 | 0.98 ± 0.04 |
| No Tail | Rearing Supported [5] | **0.93** ± **0.07** | **0.94** ± **0.09** | 0.90 ± 0.11 | **0.91** ± **0.09** | **0.98** ± **0.03** |
| No Paws | Rearing Supported [5] | 0.80 ± 0.14 | 0.79 ± 0.20 | 0.84 ± 0.15 | 0.79 ± 0.13 | 0.91 ± 0.12 |
| No Nose or Tail | Rearing Supported [5] | **0.93** ± **0.07** | 0.94 ± 0.10 | **0.90** ± **0.12** | 0.91 ± 0.10 | **0.98** ± **0.03** |
| Nose and Paws | Rearing Supported [5] | 0.92 ± 0.10 | 0.92 ± 0.12 | 0.87 ± 0.17 | 0.89 ± 0.14 | 0.96 ± 0.08 |
| Nose and Base Tail | Rearing Supported [5] | 0.74 ± 0.11 | 0.72 ± 0.16 | 0.70 ± 0.16 | 0.70 ± 0.13 | 0.82 ± 0.13 |
| [5] | Rearing Unsupported [5] | 0.83 ± 0.13 | 0.83 ± 0.24 | 0.73 ± 0.21 | 0.75 ± 0.20 | 0.91 ± 0.10 |
| [10] | Rearing Unsupported [5] | 0.78 ± 0.09 | 0.77 ± 0.19 | 0.63 ± 0.23 | 0.65 ± 0.17 | 0.88 ± 0.10 |
| [14] | Rearing Unsupported [5] | 0.80 ± 0.09 | 0.80 ± 0.15 | 0.63 ± 0.21 | 0.67 ± 0.16 | 0.89 ± 0.08 |
| [37] | Rearing Unsupported [5] | 0.78 ± 0.10 | 0.78 ± 0.16 | 0.58 ± 0.21 | 0.64 ± 0.17 | 0.85 ± 0.10 |
| No Ears | Rearing Unsupported [5] | 0.84 ± 0.10 | 0.81 ± 0.21 | 0.72 ± 0.22 | 0.74 ± 0.19 | 0.92 ± 0.09 |
| No Tail | Rearing Unsupported [5] | 0.84 ± 0.13 | 0.84 ± 0.23 | 0.73 ± 0.22 | 0.76 ± 0.20 | 0.92 ± 0.09 |
| No Paws | Rearing Unsupported [5] | 0.80 ± 0.08 | 0.82 ± 0.18 | 0.62 ± 0.25 | 0.66 ± 0.17 | 0.88 ± 0.10 |
| No Nose or Tail | Rearing [5] Unsupported | **0.85** ± **0.08** | **0.85** ± **0.17** | **0.74** ± **0.18** | **0.77** ± **0.14** | **0.93** ± **0.07** |
| Nose and Paws | Rearing [5] Unsupported | 0.82 ± 0.10 | 0.82 ± 0.21 | 0.68 ± 0.21 | 0.72 ± 0.18 | 0.91 ± 0.10 |
| Nose and Base Tail | Rearing [5] Unsupported | 0.78 ± 0.08 | 0.75 ± 0.18 | 0.60 ± 0.16 | 0.65 ± 0.13 | 0.83 ± 0.09 |
| [5] | Scratch [5] | **0.91** ± **0.05** | 0.92 ± 0.06 | 0.89 ± 0.09 | 0.90 ± 0.05 | **0.98** ± **0.02** |
| [10] | Scratch [5] | 0.90 ± 0.05 | 0.90 ± 0.07 | 0.89 ± 0.09 | 0.89 ± 0.05 | 0.97 ± 0.02 |
| [14] | Scratch [5] | 0.82 ± 0.12 | 0.88 ± 0.07 | 0.76 ± 0.18 | 0.80 ± 0.12 | 0.91 ± 0.08 |
| [37] | Scratch [5] | 0.77 ± 0.09 | 0.82 ± 0.09 | 0.71 ± 0.16 | 0.75 ± 0.11 | 0.88 ± 0.05 |
| No Ears | Scratch [5] | 0.91 ± 0.06 | **0.93** ± **0.06** | 0.89 ± 0.11 | 0.90 ± 0.06 | 0.98 ± 0.02 |
| No Tail | Scratch [5] | **0.91** ± **0.05** | 0.92 ± 0.07 | **0.90** ± **0.10** | **0.91** ± **0.06** | **0.98** ± **0.02** |
| No Paws | Scratch [5] | 0.84 ± 0.10 | 0.88 ± 0.08 | 0.81 ± 0.14 | 0.83 ± 0.09 | 0.93 ± 0.07 |
| No Nose or Tail | Scratch [5] | 0.90 ± 0.04 | 0.92 ± 0.07 | 0.89 ± 0.10 | 0.90 ± 0.05 | 0.98 ± 0.03 |
| Nose and Paws | Scratch [5] | 0.90 ± 0.06 | 0.91 ± 0.08 | 0.88 ± 0.14 | 0.89 ± 0.09 | 0.97 ± 0.04 |
| Nose and Base Tail | Scratch [5] | 0.74 ± 0.09 | 0.77 ± 0.10 | 0.70 ± 0.15 | 0.72 ± 0.08 | 0.83 ± 0.08 |
| [5] | Left Turn [5] | **0.90** ± **0.05** | 0.81 ± 0.13 | 0.74 ± 0.15 | **0.76** ± **0.10** | 0.94 ± 0.05 |
| [10] | Left Turn [5] | **0.90** ± **0.05** | **0.84** ± **0.13** | 0.71 ± 0.17 | 0.75 ± 0.12 | 0.94 ± 0.05 |
| [14] | Left Turn [5] | 0.89 ± 0.05 | 0.83 ± 0.14 | 0.70 ± 0.16 | 0.75 ± 0.12 | 0.94 ± 0.05 |
| [37] | Left Turn [5] | 0.89 ± 0.05 | 0.82 ± 0.13 | 0.69 ± 0.17 | 0.74 ± 0.13 | 0.93 ± 0.05 |
| No Ears | Left Turn [5] | 0.89 ± 0.04 | **0.84** ± **0.13** | 0.70 ± 0.13 | 0.75 ± 0.09 | 0.94 ± 0.05 |
| No Tail | Left Turn [5] | **0.90** ±<br>**0.05** | 0.81 ±<br>0.12 | **0.75** ±<br>**0.17** | 0.76 ±<br>0.11 | **0.95** ±<br>**0.05** |
| No Paws | Left Turn [5] | **0.90** ±<br>**0.05** | 0.84 ±<br>0.15 | 0.72 ±<br>0.13 | **0.76** ±<br>**0.10** | 0.94 ±<br>0.05 |
| No Nose<br>or Tail | Left Turn [5] | 0.88 ±<br>0.05 | 0.81 ±<br>0.14 | 0.67 ±<br>0.17 | 0.72 ±<br>0.13 | 0.93 ±<br>0.05 |
| Nose and<br>Paws | Left Turn [5] | 0.88 ±<br>0.06 | 0.82 ±<br>0.16 | 0.69 ±<br>0.17 | 0.72 ±<br>0.14 | 0.93 ±<br>0.05 |
| Nose and<br>Base Tail | Left Turn [5] | 0.88 ±<br>0.06 | 0.79 ±<br>0.14 | 0.71 ±<br>0.16 | 0.73 ±<br>0.13 | 0.92 ±<br>0.05 |
| [5] | Right Turn [5] | **0.92** ±<br>**0.05** | 0.84 ±<br>0.15 | **0.91** ±<br>**0.08** | 0.86 ±<br>0.09 | 0.97 ±<br>0.03 |
| [10] | Right Turn [5] | 0.91 ±<br>0.06 | 0.85 ±<br>0.14 | 0.81 ±<br>0.17 | 0.82 ±<br>0.11 | 0.96 ±<br>0.04 |
| [14] | Right Turn [5] | 0.91 ±<br>0.05 | **0.86** ±<br>**0.12** | 0.84 ±<br>0.13 | 0.84 ±<br>0.08 | **0.97** ±<br>**0.02** |
| [37] | Right Turn [5] | 0.90 ±<br>0.07 | 0.85 ±<br>0.11 | 0.83 ±<br>0.14 | 0.83 ±<br>0.09 | **0.97** ±<br>**0.02** |
| No Ears | Right Turn [5] | 0.91 ±<br>0.05 | 0.85 ±<br>0.14 | 0.86 ±<br>0.10 | 0.85 ±<br>0.08 | 0.97 ±<br>0.03 |
| No Tail | Right Turn [5] | **0.92** ±<br>**0.05** | 0.85 ±<br>0.14 | **0.91** ±<br>**0.08** | **0.87** ±<br>**0.08** | 0.97 ±<br>0.03 |
| No Paws | Right Turn [5] | 0.91 ±<br>0.05 | 0.85 ±<br>0.12 | 0.83 ±<br>0.15 | 0.83 ±<br>0.10 | 0.97 ±<br>0.03 |
| No Nose<br>or Tail | Right Turn [5] | 0.91 ±<br>0.05 | 0.84 ±<br>0.12 | 0.85 ±<br>0.12 | 0.84 ±<br>0.08 | 0.97 ±<br>0.03 |
| Nose and<br>Paws | Right Turn [5] | 0.89 ±<br>0.06 | 0.82 ±<br>0.13 | 0.83 ±<br>0.12 | 0.81 ±<br>0.08 | 0.96 ±<br>0.04 |
| Nose and<br>Base Tail | Right Turn [5] | 0.89 ±<br>0.07 | 0.81 ±<br>0.12 | 0.83 ±<br>0.13 | 0.81 ±<br>0.10 | 0.96 ±<br>0.04 |
| [5] | Rearing Supported [4] | 0.95 ±<br>0.01 | 0.91 ±<br>0.04 | 0.82 ±<br>0.05 | 0.86 ±<br>0.03 | **0.98** ±<br>**0.01** |
| [10] | Rearing Supported [4] | 0.94 ±<br>0.01 | 0.89 ±<br>0.06 | 0.78 ±<br>0.07 | 0.83 ±<br>0.03 | 0.97 ±<br>0.01 |
| [14] | Rearing Supported [4] | 0.94 ±<br>0.02 | 0.88 ±<br>0.05 | 0.76 ±<br>0.06 | 0.81 ±<br>0.03 | 0.97 ±<br>0.01 |
| [37] | Rearing Supported [4] | 0.94 ± 0.02 | 0.87 ± 0.06 | 0.75 ± 0.06 | 0.81 ± 0.03 | 0.97 ± 0.01 |
| No Ears | Rearing Supported [4] | 0.94 ± 0.02 | 0.89 ± 0.06 | 0.78 ± 0.08 | 0.83 ± 0.03 | **0.98 ± 0.01** |
| No Tail | Rearing Supported [4] | **0.96 ± 0.01** | **0.91 ± 0.03** | **0.83 ± 0.05** | **0.87 ± 0.03** | **0.98 ± 0.01** |
| No Paws | Rearing Supported [4] | 0.94 ± 0.02 | 0.88 ± 0.05 | 0.75 ± 0.07 | 0.81 ± 0.03 | 0.97 ± 0.01 |
| No Nose or Tail | Rearing Supported [4] | 0.93 ± 0.02 | 0.87 ± 0.05 | 0.72 ± 0.07 | 0.79 ± 0.03 | 0.96 ± 0.01 |
| Nose and Paws | Rearing Supported [4] | 0.91 ± 0.02 | 0.82 ± 0.08 | 0.65 ± 0.10 | 0.71 ± 0.05 | 0.94 ± 0.02 |
| Nose and Base Tail | Rearing Supported [4] | 0.89 ± 0.02 | 0.76 ± 0.07 | 0.54 ± 0.08 | 0.63 ± 0.05 | 0.90 ± 0.02 |
| [5] | Rearing Unsupported [4] | **0.93 ± 0.03** | 0.71 ± 0.20 | **0.38 ± 0.11** | **0.49 ± 0.14** | 0.90 ± 0.03 |
| [10] | Rearing Unsupported [4] | 0.92 ± 0.03 | 0.68 ± 0.20 | 0.34 ± 0.10 | 0.45 ± 0.13 | 0.90 ± 0.03 |
| [14] | Rearing Unsupported [4] | 0.92 ± 0.03 | 0.66 ± 0.21 | 0.33 ± 0.10 | 0.44 ± 0.13 | 0.89 ± 0.04 |
| [37] | Rearing Unsupported [4] | 0.92 ± 0.03 | 0.65 ± 0.20 | 0.33 ± 0.11 | 0.43 ± 0.13 | 0.89 ± 0.03 |
| No Ears | Rearing Unsupported [4] | 0.92 ± 0.03 | 0.68 ± 0.20 | 0.35 ± 0.10 | 0.45 ± 0.13 | 0.90 ± 0.03 |
| No Tail | Rearing Unsupported [4] | **0.93 ± 0.03** | **0.71 ± 0.19** | 0.37 ± 0.10 | 0.48 ± 0.13 | **0.91 ± 0.03** |
| No Paws | Rearing Unsupported [4] | 0.92 ± 0.03 | 0.66 ± 0.20 | 0.34 ± 0.11 | 0.44 ± 0.14 | 0.89 ± 0.03 |
| No Nose or Tail | Rearing Unsupported [4] | 0.92 ± 0.03 | 0.65 ± 0.20 | 0.31 ± 0.09 | 0.41 ± 0.12 | 0.87 ± 0.03 |
| Nose and Paws | Rearing Unsupported [4] | 0.91 ± 0.03 | 0.62 ± 0.22 | 0.26 ± 0.11 | 0.36 ± 0.14 | 0.86 ± 0.03 |
| Nose and Base Tail | Rearing Unsupported [4] | 0.90 ± 0.04 | 0.49 ± 0.21 | 0.16 ± 0.09 | 0.23 ± 0.11 | 0.82 ± 0.03 |

**Table S4:**
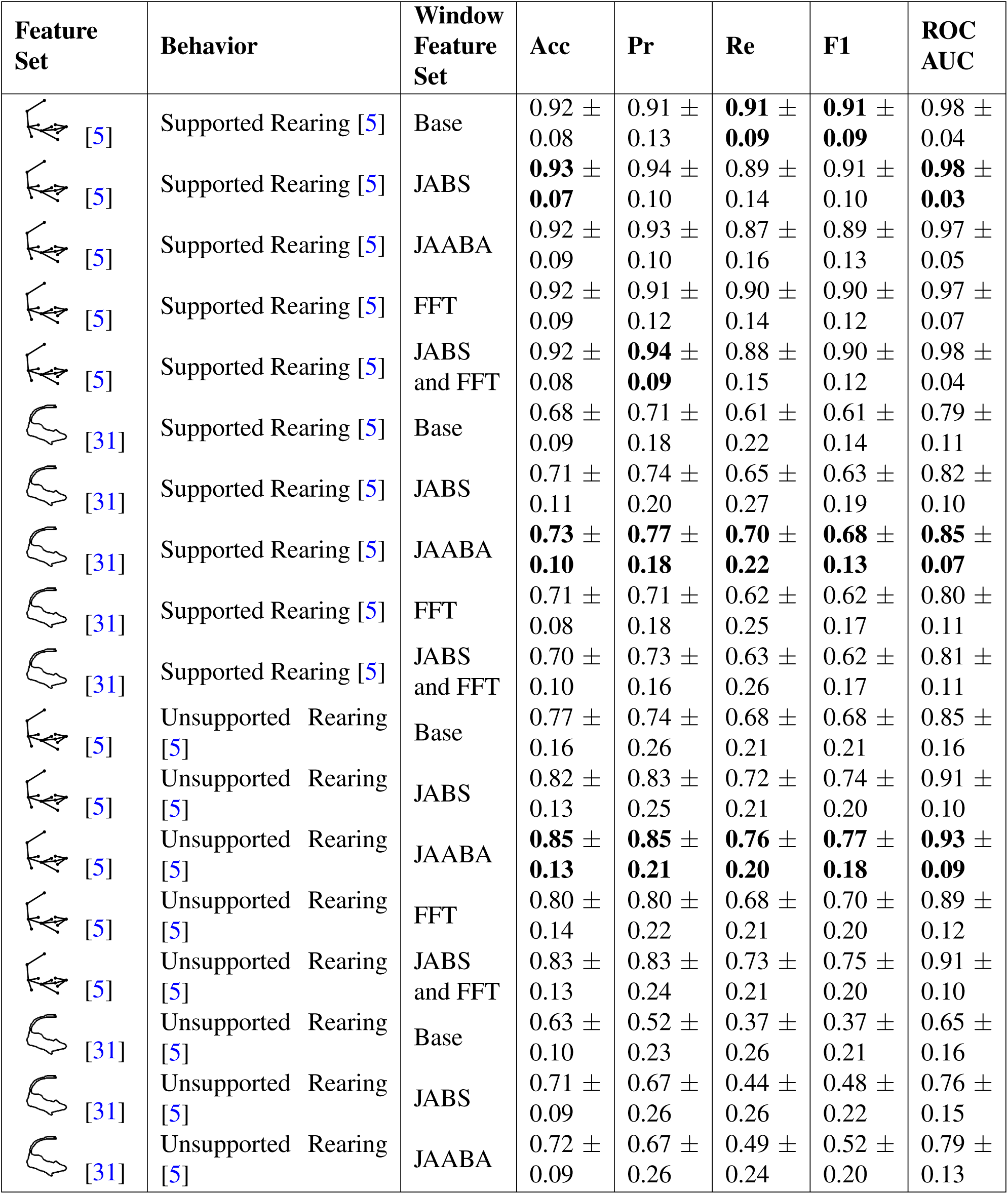
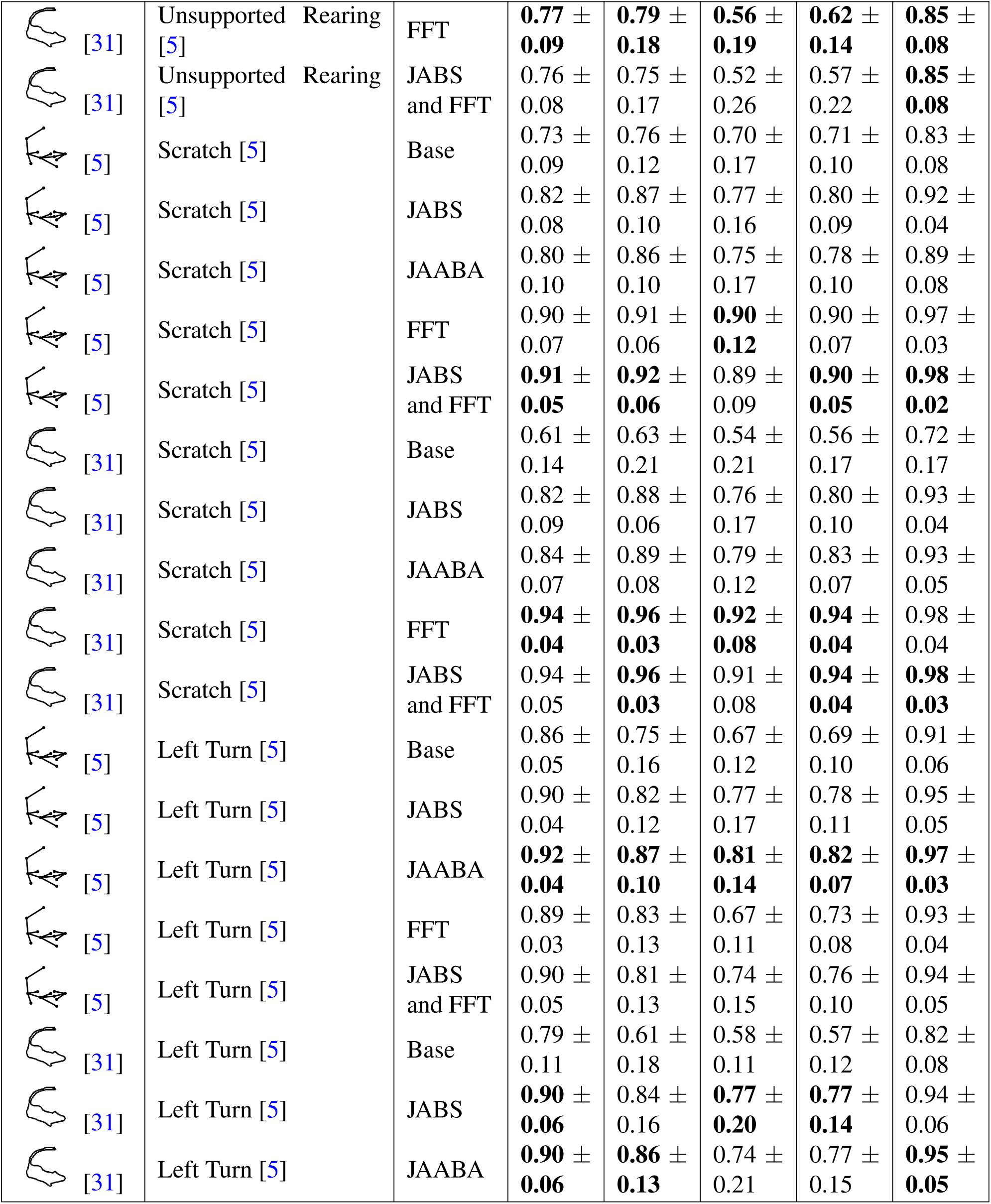
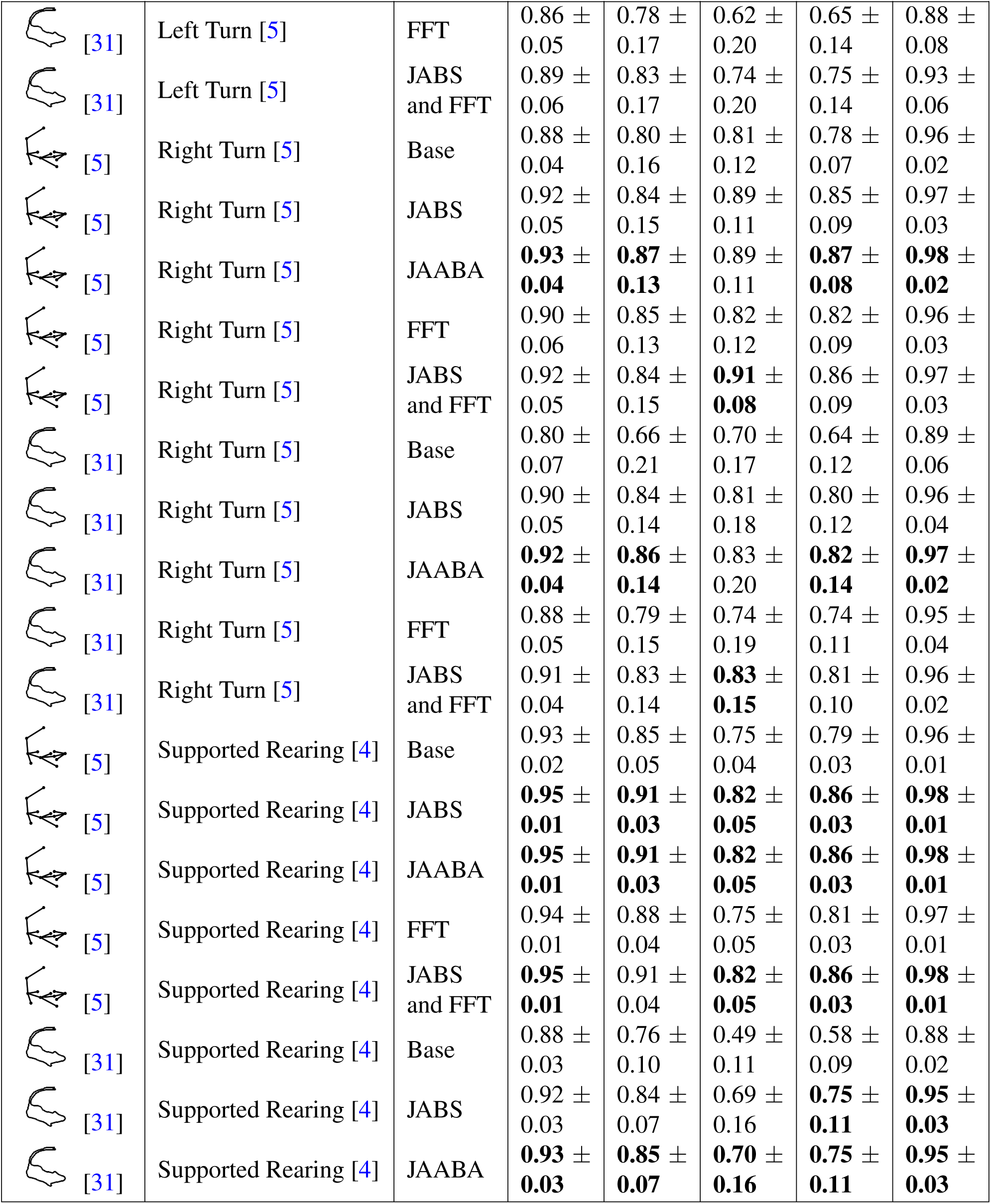
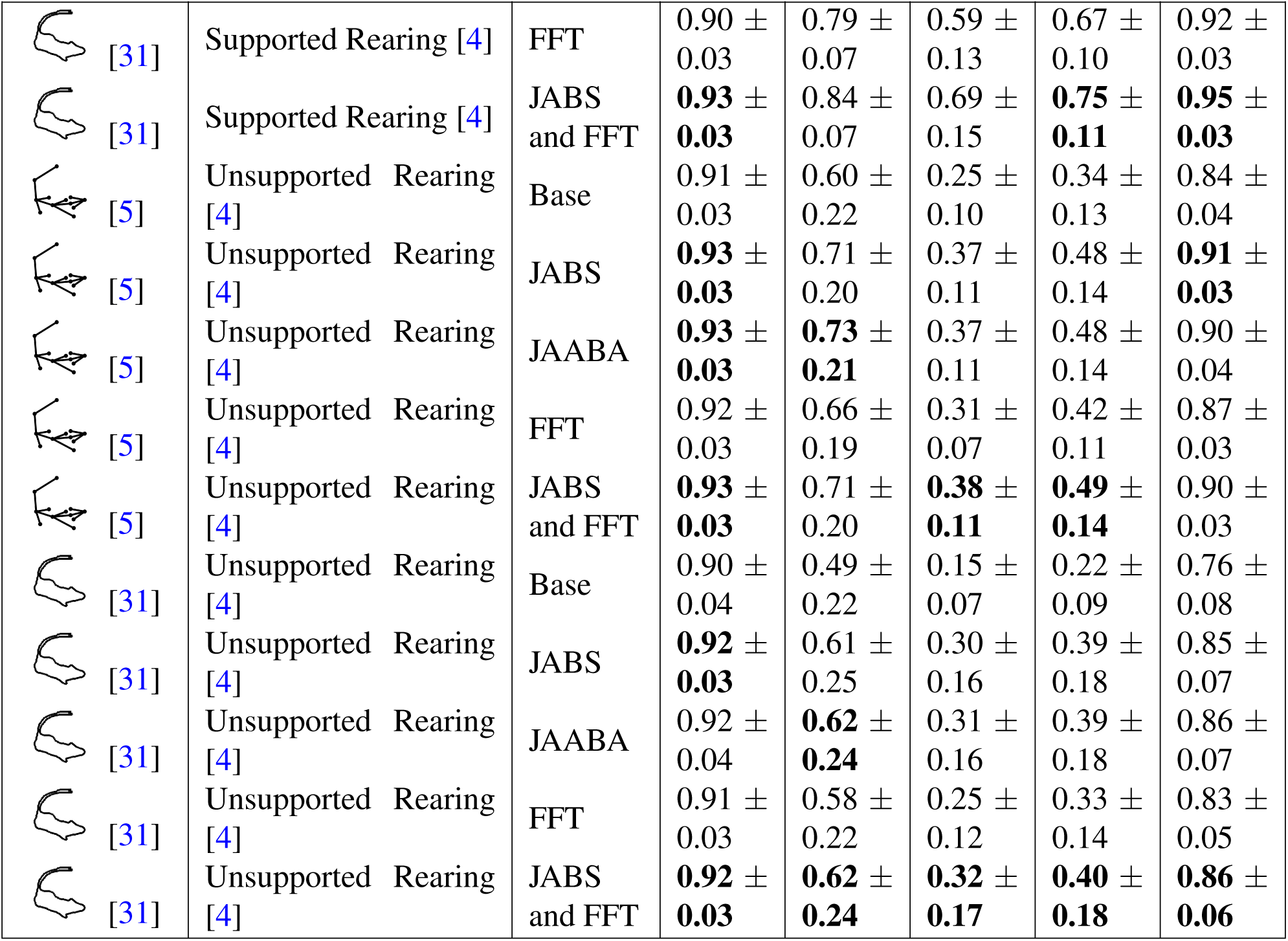
Performance of classification task (mean *±* std) of additional behaviors for segmentation and keypoint representations with different window features. Bold indicates best performance.

| Feature Set | Behavior | Window Feature Set | Acc | Pr | Re | F1 | ROC AUC |
| --- | --- | --- | --- | --- | --- | --- | --- |
| [5] | Supported Rearing [5] | Base | 0.92 ± 0.08 | 0.91 ± 0.13 | **0.91** ± **0.09** | **0.91** ± **0.09** | 0.98 ± 0.04 |
| [5] | Supported Rearing [5] | JABS | **0.93** ± **0.07** | 0.94 ± 0.10 | 0.89 ± 0.14 | 0.91 ± 0.10 | **0.98** ± **0.03** |
| [5] | Supported Rearing [5] | JAABA | 0.92 ± 0.09 | 0.93 ± 0.10 | 0.87 ± 0.16 | 0.89 ± 0.13 | 0.97 ± 0.05 |
| [5] | Supported Rearing [5] | FFT | 0.92 ± 0.09 | 0.91 ± 0.12 | 0.90 ± 0.14 | 0.90 ± 0.12 | 0.97 ± 0.07 |
| [5] | Supported Rearing [5] | JABS and FFT | 0.92 ± 0.08 | **0.94** ± **0.09** | 0.88 ± 0.15 | 0.90 ± 0.12 | 0.98 ± 0.04 |
| [31] | Supported Rearing [5] | Base | 0.68 ± 0.09 | 0.71 ± 0.18 | 0.61 ± 0.22 | 0.61 ± 0.14 | 0.79 ± 0.11 |
| [31] | Supported Rearing [5] | JABS | 0.71 ± 0.11 | 0.74 ± 0.20 | 0.65 ± 0.27 | 0.63 ± 0.19 | 0.82 ± 0.10 |
| [31] | Supported Rearing [5] | JAABA | **0.73** ± **0.10** | **0.77** ± **0.18** | **0.70** ± **0.22** | **0.68** ± **0.13** | **0.85** ± **0.07** |
| [31] | Supported Rearing [5] | FFT | 0.71 ± 0.08 | 0.71 ± 0.18 | 0.62 ± 0.25 | 0.62 ± 0.17 | 0.80 ± 0.11 |
| [31] | Supported Rearing [5] | JABS and FFT | 0.70 ± 0.10 | 0.73 ± 0.16 | 0.63 ± 0.26 | 0.62 ± 0.17 | 0.81 ± 0.11 |
| [5] | Unsupported Rearing [5] | Base | 0.77 ± 0.16 | 0.74 ± 0.26 | 0.68 ± 0.21 | 0.68 ± 0.21 | 0.85 ± 0.16 |
| [5] | Unsupported Rearing [5] | JABS | 0.82 ± 0.13 | 0.83 ± 0.25 | 0.72 ± 0.21 | 0.74 ± 0.20 | 0.91 ± 0.10 |
| [5] | Unsupported Rearing [5] | JAABA | **0.85** ± **0.13** | **0.85** ± **0.21** | **0.76** ± **0.20** | **0.77** ± **0.18** | **0.93** ± **0.09** |
| [5] | Unsupported Rearing [5] | FFT | 0.80 ± 0.14 | 0.80 ± 0.22 | 0.68 ± 0.21 | 0.70 ± 0.20 | 0.89 ± 0.12 |
| [5] | Unsupported Rearing [5] | JABS and FFT | 0.83 ± 0.13 | 0.83 ± 0.24 | 0.73 ± 0.21 | 0.75 ± 0.20 | 0.91 ± 0.10 |
| [31] | Unsupported Rearing [5] | Base | 0.63 ± 0.10 | 0.52 ± 0.23 | 0.37 ± 0.26 | 0.37 ± 0.21 | 0.65 ± 0.16 |
| [31] | Unsupported Rearing [5] | JABS | 0.71 ± 0.09 | 0.67 ± 0.26 | 0.44 ± 0.26 | 0.48 ± 0.22 | 0.76 ± 0.15 |
| [31] | Unsupported Rearing [5] | JAABA | 0.72 ± 0.09 | 0.67 ± 0.26 | 0.49 ± 0.24 | 0.52 ± 0.20 | 0.79 ± 0.13 |

| Feature Set | Behavior | Window Features | Acc | Pr | Re | F1 | ROC AUC |
| --- | --- | --- | --- | --- | --- | --- | --- |
| [31] | Unsupported Rearing [5] | FFT | **0.77** ± **0.09** | **0.79** ± **0.18** | **0.56** ± **0.19** | **0.62** ± **0.14** | **0.85** ± **0.08** |
| [31] | Unsupported Rearing [5] | JABS and FFT | 0.76 ± 0.08 | 0.75 ± 0.17 | 0.52 ± 0.26 | 0.57 ± 0.22 | **0.85** ± **0.08** |
| [5] | Scratch [5] | Base | 0.73 ± 0.09 | 0.76 ± 0.12 | 0.70 ± 0.17 | 0.71 ± 0.10 | 0.83 ± 0.08 |
| [5] | Scratch [5] | JABS | 0.82 ± 0.08 | 0.87 ± 0.10 | 0.77 ± 0.16 | 0.80 ± 0.09 | 0.92 ± 0.04 |
| [5] | Scratch [5] | JAABA | 0.80 ± 0.10 | 0.86 ± 0.10 | 0.75 ± 0.17 | 0.78 ± 0.10 | 0.89 ± 0.08 |
| [5] | Scratch [5] | FFT | 0.90 ± 0.07 | 0.91 ± 0.06 | **0.90** ± **0.12** | 0.90 ± 0.07 | 0.97 ± 0.03 |
| [5] | Scratch [5] | JABS and FFT | **0.91** ± **0.05** | **0.92** ± **0.06** | 0.89 ± 0.09 | **0.90** ± **0.05** | **0.98** ± **0.02** |
| [31] | Scratch [5] | Base | 0.61 ± 0.14 | 0.63 ± 0.21 | 0.54 ± 0.21 | 0.56 ± 0.17 | 0.72 ± 0.17 |
| [31] | Scratch [5] | JABS | 0.82 ± 0.09 | 0.88 ± 0.06 | 0.76 ± 0.17 | 0.80 ± 0.10 | 0.93 ± 0.04 |
| [31] | Scratch [5] | JAABA | 0.84 ± 0.07 | 0.89 ± 0.08 | 0.79 ± 0.12 | 0.83 ± 0.07 | 0.93 ± 0.05 |
| [31] | Scratch [5] | FFT | **0.94** ± **0.04** | **0.96** ± **0.03** | **0.92** ± **0.08** | **0.94** ± **0.04** | 0.98 ± 0.04 |
| [31] | Scratch [5] | JABS and FFT | 0.94 ± 0.05 | **0.96** ± **0.03** | 0.91 ± 0.08 | **0.94** ± **0.04** | **0.98** ± **0.03** |
| [5] | Left Turn [5] | Base | 0.86 ± 0.05 | 0.75 ± 0.16 | 0.67 ± 0.12 | 0.69 ± 0.10 | 0.91 ± 0.06 |
| [5] | Left Turn [5] | JABS | 0.90 ± 0.04 | 0.82 ± 0.12 | 0.77 ± 0.17 | 0.78 ± 0.11 | 0.95 ± 0.05 |
| [5] | Left Turn [5] | JAABA | **0.92** ± **0.04** | **0.87** ± **0.10** | **0.81** ± **0.14** | **0.82** ± **0.07** | **0.97** ± **0.03** |
| [5] | Left Turn [5] | FFT | 0.89 ± 0.03 | 0.83 ± 0.13 | 0.67 ± 0.11 | 0.73 ± 0.08 | 0.93 ± 0.04 |
| [5] | Left Turn [5] | JABS and FFT | 0.90 ± 0.05 | 0.81 ± 0.13 | 0.74 ± 0.15 | 0.76 ± 0.10 | 0.94 ± 0.05 |
| [31] | Left Turn [5] | Base | 0.79 ± 0.11 | 0.61 ± 0.18 | 0.58 ± 0.11 | 0.57 ± 0.12 | 0.82 ± 0.08 |
| [31] | Left Turn [5] | JABS | **0.90** ± **0.06** | 0.84 ± 0.16 | **0.77** ± **0.20** | **0.77** ± **0.14** | 0.94 ± 0.06 |
| [31] | Left Turn [5] | JAABA | **0.90** ± **0.06** | **0.86** ± **0.13** | 0.74 ± 0.21 | 0.77 ± 0.15 | **0.95** ± **0.05** |
| [31] | Left Turn [5] | FFT | 0.86 ± 0.05 | 0.78 ± 0.17 | 0.62 ± 0.20 | 0.65 ± 0.14 | 0.88 ± 0.08 |
| [31] | Left Turn [5] | JABS and FFT | 0.89 ± 0.06 | 0.83 ± 0.17 | 0.74 ± 0.20 | 0.75 ± 0.14 | 0.93 ± 0.06 |
| [5] | Right Turn [5] | Base | 0.88 ± 0.04 | 0.80 ± 0.16 | 0.81 ± 0.12 | 0.78 ± 0.07 | 0.96 ± 0.02 |
| [5] | Right Turn [5] | JABS | 0.92 ± 0.05 | 0.84 ± 0.15 | 0.89 ± 0.11 | 0.85 ± 0.09 | 0.97 ± 0.03 |
| [5] | Right Turn [5] | JAABA | **0.93 ± 0.04** | **0.87 ± 0.13** | 0.89 ± 0.11 | **0.87 ± 0.08** | **0.98 ± 0.02** |
| [5] | Right Turn [5] | FFT | 0.90 ± 0.06 | 0.85 ± 0.13 | 0.82 ± 0.12 | 0.82 ± 0.09 | 0.96 ± 0.03 |
| [5] | Right Turn [5] | JABS and FFT | 0.92 ± 0.05 | 0.84 ± 0.15 | **0.91 ± 0.08** | 0.86 ± 0.09 | 0.97 ± 0.03 |
| [31] | Right Turn [5] | Base | 0.80 ± 0.07 | 0.66 ± 0.21 | 0.70 ± 0.17 | 0.64 ± 0.12 | 0.89 ± 0.06 |
| [31] | Right Turn [5] | JABS | 0.90 ± 0.05 | 0.84 ± 0.14 | 0.81 ± 0.18 | 0.80 ± 0.12 | 0.96 ± 0.04 |
| [31] | Right Turn [5] | JAABA | **0.92 ± 0.04** | **0.86 ± 0.14** | 0.83 ± 0.20 | **0.82 ± 0.14** | **0.97 ± 0.02** |
| [31] | Right Turn [5] | FFT | 0.88 ± 0.05 | 0.79 ± 0.15 | 0.74 ± 0.19 | 0.74 ± 0.11 | 0.95 ± 0.04 |
| [31] | Right Turn [5] | JABS and FFT | 0.91 ± 0.04 | 0.83 ± 0.14 | **0.83 ± 0.15** | 0.81 ± 0.10 | 0.96 ± 0.02 |
| [5] | Supported Rearing [4] | Base | 0.93 ± 0.02 | 0.85 ± 0.05 | 0.75 ± 0.04 | 0.79 ± 0.03 | 0.96 ± 0.01 |
| [5] | Supported Rearing [4] | JABS | **0.95 ± 0.01** | **0.91 ± 0.03** | **0.82 ± 0.05** | **0.86 ± 0.03** | **0.98 ± 0.01** |
| [5] | Supported Rearing [4] | JAABA | **0.95 ± 0.01** | **0.91 ± 0.03** | **0.82 ± 0.05** | **0.86 ± 0.03** | **0.98 ± 0.01** |
| [5] | Supported Rearing [4] | FFT | 0.94 ± 0.01 | 0.88 ± 0.04 | 0.75 ± 0.05 | 0.81 ± 0.03 | 0.97 ± 0.01 |
| [5] | Supported Rearing [4] | JABS and FFT | **0.95 ± 0.01** | 0.91 ± 0.04 | **0.82 ± 0.05** | **0.86 ± 0.03** | **0.98 ± 0.01** |
| [31] | Supported Rearing [4] | Base | 0.88 ± 0.03 | 0.76 ± 0.10 | 0.49 ± 0.11 | 0.58 ± 0.09 | 0.88 ± 0.02 |
| [31] | Supported Rearing [4] | JABS | 0.92 ± 0.03 | 0.84 ± 0.07 | 0.69 ± 0.16 | **0.75 ± 0.11** | **0.95 ± 0.03** |
| [31] | Supported Rearing [4] | JAABA | **0.93 ± 0.03** | **0.85 ± 0.07** | **0.70 ± 0.16** | **0.75 ± 0.11** | **0.95 ± 0.03** |
| [31] | Supported Rearing [4] | FFT | 0.90 ± 0.03 | 0.79 ± 0.07 | 0.59 ± 0.13 | 0.67 ± 0.10 | 0.92 ± 0.03 |
| [31] | Supported Rearing [4] | JABS and FFT | **0.93 ± 0.03** | 0.84 ± 0.07 | 0.69 ± 0.15 | **0.75 ± 0.11** | **0.95 ± 0.03** |
| [5] | Unsupported Rearing [4] | Base | 0.91 ± 0.03 | 0.60 ± 0.22 | 0.25 ± 0.10 | 0.34 ± 0.13 | 0.84 ± 0.04 |
| [5] | Unsupported Rearing [4] | JABS | **0.93 ± 0.03** | 0.71 ± 0.20 | 0.37 ± 0.11 | 0.48 ± 0.14 | **0.91 ± 0.03** |
| [5] | Unsupported Rearing [4] | JAABA | **0.93 ± 0.03** | **0.73 ± 0.21** | 0.37 ± 0.11 | 0.48 ± 0.14 | 0.90 ± 0.04 |
| [5] | Unsupported Rearing [4] | FFT | 0.92 ± 0.03 | 0.66 ± 0.19 | 0.31 ± 0.07 | 0.42 ± 0.11 | 0.87 ± 0.03 |
| [5] | Unsupported Rearing [4] | JABS and FFT | **0.93 ± 0.03** | 0.71 ± 0.20 | **0.38 ± 0.11** | **0.49 ± 0.14** | 0.90 ± 0.03 |
| [31] | Unsupported Rearing [4] | Base | 0.90 ± 0.04 | 0.49 ± 0.22 | 0.15 ± 0.07 | 0.22 ± 0.09 | 0.76 ± 0.08 |
| [31] | Unsupported Rearing [4] | JABS | **0.92 ± 0.03** | 0.61 ± 0.25 | 0.30 ± 0.16 | 0.39 ± 0.18 | 0.85 ± 0.07 |
| [31] | Unsupported Rearing [4] | JAABA | 0.92 ± 0.04 | **0.62 ± 0.24** | 0.31 ± 0.16 | 0.39 ± 0.18 | 0.86 ± 0.07 |
| [31] | Unsupported Rearing [4] | FFT | 0.91 ± 0.03 | 0.58 ± 0.22 | 0.25 ± 0.12 | 0.33 ± 0.14 | 0.83 ± 0.05 |
| [31] | Unsupported Rearing [4] | JABS and FFT | **0.92 ± 0.03** | **0.62 ± 0.24** | **0.32 ± 0.17** | **0.40 ± 0.18** | **0.86 ± 0.06** |

## S1 Methods

### S1.1 Behavioral Datasets

We run our experiments using a combination of our publicly available annotated datasets (JABS Supported Rearing, JABS Unsupported Rearing, Scratch, Left Turn, Right Turn, Grooming) [5, 38], and other datasets made available by Sturman et al. (Sturman Supported Rearing and Sturman Unsupported Rearing) [4] (Table S6). While these are not the only behavioral datasets made public for conducting these tests, we selected them since we provide the models for our datasets that operate directly on raw frame data to produce segmentation predictions [3] and keypoint predictions [29]. Sturman et al. [4] released their internal predicted keypoints on their datasets as well as behavioral annotations. We use SAM2 [33] to generate segmentation features for the Sturman dataset, as it is published without any segmentation. Having raw frame inputs allows both pose and segmentation based feature abstraction tests to be conducted. We also use the grooming dataset [38] for scaling experiments since it is the largest annotated lab dataset of single animals currently released for conducting feature optimization experiments.

### S1.2 Keypoint Features

#### Keypoint Prediction Model

We used two different methods for acquiring keypoint predictions. First, for the behavior datasets JABS Supported Rearing [5], JABS Unsupported Rearing [5], Scratch, Left Turn, Right Turn, and Grooming behaviors, we use our previously trained keypoint model [29]. We inferred our trained 12-keypoint prediction model on the raw video frames. We then stored keypoint prediction locations for every frame such that features could be calculated from them. For the Sturman datasets we accessed the precomputed keypoint predictions made available in [4]. We used these predictions in our downstream analysis and calculated features consistently across all datasets.

#### Keypoint Feature Descriptions

We adopted the keypoint feature set described in [5], which includes pairwise distances, three keypoint angles, and individual keypoint velocity terms. In total, there are 110 keypoint-based features described for the 12-keypoint model. Since Sturman et al. [4] includes 13 keypoints, we mapped and subsetted those keypoints to match the 12-keypoint representation from Choudhary et al. [5] for consistency.

### S1.3 Segmentation Feature

#### Segmentation Prediction Model

For behavior datasets originating from our lab [38, 5], we used our trained segmentation model [3]. This model was trained to operate on visually diverse mice in an open field. Since our group both trained this model and also generated the behavior datasets, the inputs would not suffer from a distribution shift. Each frame was predicted using the model and a list of contours was subsequently stored to describe the resulting segmentation mask.

For the out-of-distribution behavior datasets in Sturman et al.[4], we adopted SAM2 [33] to predict segmentation masks using a single keypoint prompt per video. For each of the 20 videos in the behavior dataset, we prompted the center frame of the video using the center spine keypoint within the published keypoint predictions. We then propagated the prediction forward and backward in time to obtain segmentation predictions on the entire video. We also attempted SAM [32], but this approach could not adequately scale on the behavior datasets. Unlike SAM2 which could perform reasonably with 1 prompt for an entire video, SAM began making incorrect predictions as the prediction frame got further from the prompt frame. While it is possible to prompt at a higher frequency, this cost would prohibit adoption by behavioral lab groups for large scale experiments.

#### Segmentation Feature Descriptions

Earlier experiments in the field focused on using ellipse approximation-based segmentation features for training classifiers. Here, we adopt segmentation features from the field of shape descriptors [48, 54, 53]. Using a similar approach to the egocentric design of keypoint features, we select segmentation descriptors which are translation invariant. We do not control for the orientation of the mouse, and do not restrict features to be rotational invariant. In total, we selected 32 base segmentation features. See Supplemental Section S0.3 for a full list of segmentation features.

### S1.4 Temporal Feature Design

The inclusion of temporal information into the feature vector for classification varies across different applications. Here we test 3 distinct approaches for the inclusion of temporal information (Table S5).

We hold the window radius constant at 16 in our approach for integrating time information. This means we allow calculations to use information from the 16 frames on either side (*t-16* to *t+16*: 33 total frames). The FFT features in [31] use a 8-15Hz band, which requires at least 30 frames. Though experiments in [5] suggest that this window size can be tuned for specific behaviors, it has shown little effect on classifier performance. We adopt signal processing features described in [31].

**Table S5:**
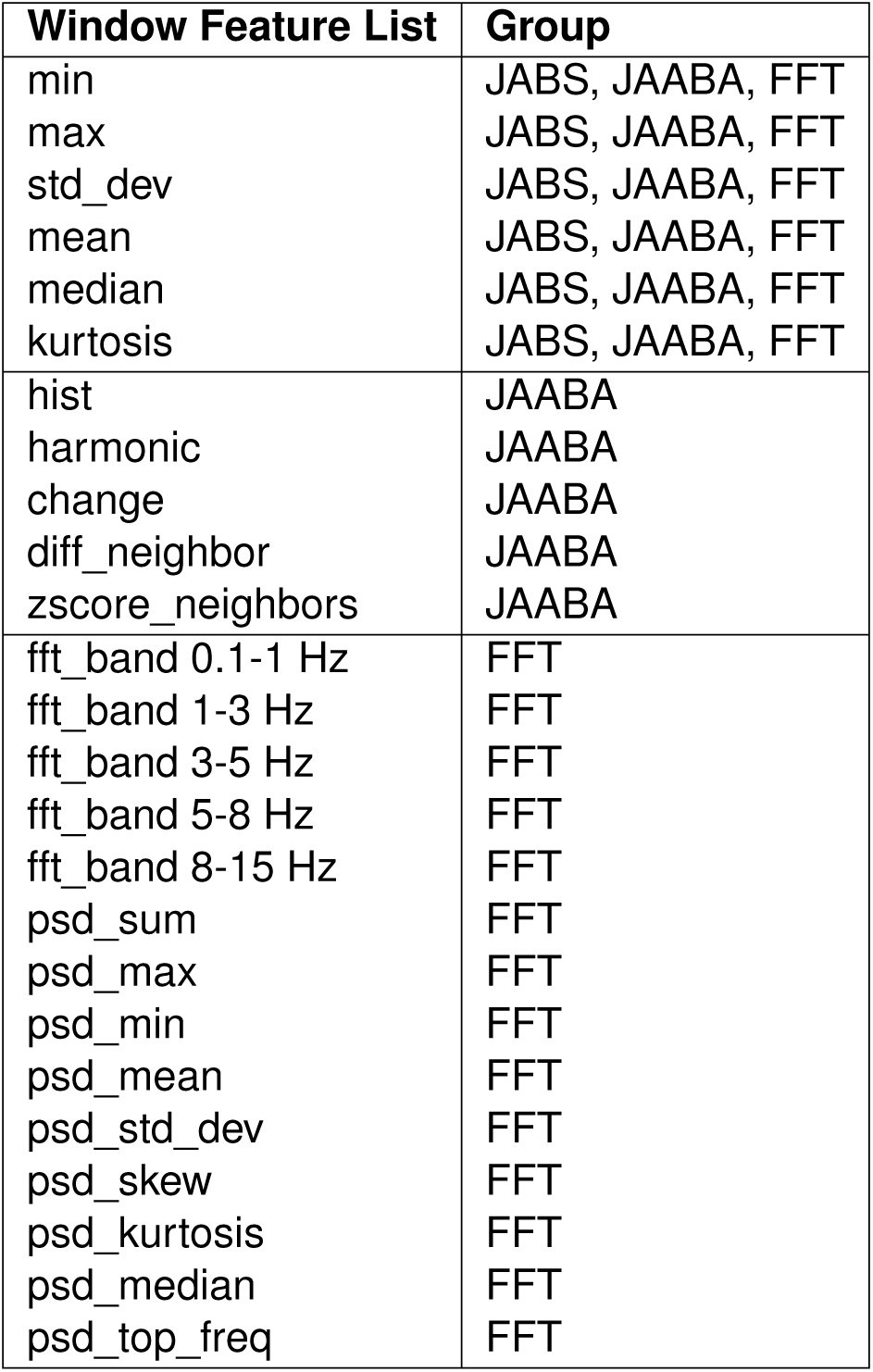
Table of window features describing implementations for JABS, JAABA, and FFT. The summary statistics present in JABS, JAABA, and FFT groups are applied to most other existing base features.

| Window Feature List | Group |
| --- | --- |
| min | JABS, JAABA, FFT |
| max | JABS, JAABA, FFT |
| std_dev | JABS, JAABA, FFT |
| mean | JABS, JAABA, FFT |
| median | JABS, JAABA, FFT |
| kurtosis | JABS, JAABA, FFT |
| hist | JAABA |
| harmonic | JAABA |
| change | JAABA |
| diff_neighbor | JAABA |
| zscore_neighbors | JAABA |
| fft_band 0.1-1 Hz | FFT |
| fft_band 1-3 Hz | FFT |
| fft_band 3-5 Hz | FFT |
| fft_band 5-8 Hz | FFT |
| fft_band 8-15 Hz | FFT |
| psd_sum | FFT |
| psd_max | FFT |
| psd_min | FFT |
| psd_mean | FFT |
| psd_std_dev | FFT |
| psd_skew | FFT |
| psd_kurtosis | FFT |
| psd_median | FFT |
| psd_top_freq | FFT |

#### JABS Window features

We adopt window features described in [5]. Briefly, these include simple summary statistics such as mean, min, max, median, and std. The feature sizes are 224 and 674 for segmentation and the full keypoints set, respectively.

#### JAABA Window Features

We adopt the window features described in [30]. Briefly, these include some summary statistics and some signal processing descriptors. The feature sizes are 448 and 1228 for segmentation and the full keypoints set, respectively.

#### FFT Window Features

We adopt signal processing features described in [31]. These features were successfully designed to be sensitive enough to detect differences in sleep states. While the original implementation uses 10s epochs, we implement the approach to use the same window radius of 16 as above. These features include more complex signal processing descriptors, such as frequency bands’ mean power spectral density (psd). The feature sizes are 672 and 1878 for segmentation and the full keypoints set, respectively.

### S1.5 Experiment Setup

#### S1.5.1 Classifier Training and Prediction

For each feature set and each behavior dataset, we train a binary XGBoost classifier to predict behavior. We store the random seed selected for training to ensure reproducible results.

#### S1.5.2 Dataset Scaling

The grooming dataset from [38] contains a large volume of annotated behavior data (2,181,790 frames). To test the influence of annotated dataset sizes, we down-sample by removing entire videos from the training set. While this approach is not optimal for training the best performing classifiers, it more closely approximates small real-world annotated dataset conditions. Generally, the lab group will annotate behavior bouts in video clips instead of single frames separated across many videos. This is the same strategy described in [38].

#### S1.5.3 Evaluation Protocol

We report frame-wise comparisons with ground truth annotations. Each behavior is analyzed and evaluated in isolation as separate datasets. For grooming behavior, the same held-out test set was used as mentioned in [38]. For all other behaviors, we use a leave-one-out cross-validation strategy. We report precision, recall, and f1-score metrics [55] along with their corresponding means and standard deviations across the cross-validation folds. While [38] describes performance using a rolling average over 46 frames of predictions to improve model performance, we report performance without any post-processing of raw predictions. When comparing across different groups, we use the same training data splits.

#### S1.5.4 Sizes and availability of real-world datasets

We present a table of behavioral annotation datasets included in our paper Table S6. To better understand the scale of real-world animal behavior sets, we find that [30] describes 19 classifiers that contain a mean annotation size of 6200 frames. We observe in literature that most lab animal behavior classifiers have fewer than 100,000 (1*e*5) frames annotated.

**Table S6:** Annotated dataset sizes across behaviors. Number of expert-labeled frames available for each behavior used in feature set evaluation. Dataset sizes reflect the practical constraints of obtaining high-quality behavioral annotations.

| **Dataset** | **Labeled Frames** |
| --- | --- |
| JABS Supported Rearing [5] | 6782 |
| JABS Unsupported Rearing [5] | 9379 |
| Scratch [5] | 9758 |
| Left turn [5] | 10034 |
| Right turn [5] | 8971 |
| Sturman Supported Rearing [4] | 304754 |
| Sturman Unsupported Rearing [4] | 304754 |
| Grooming [38] | 2181790 |

